# Phylogeographic breaks and how to find them: An empirical attempt at separating vicariance from isolation by distance in a lizard with restricted dispersal

**DOI:** 10.1101/2022.09.30.510256

**Authors:** Loïs Rancilhac, Aurélien Miralles, Philippe Geniez, Daniel Mendez-Aranda, Menad Beddek, José Carlos Brito, Raphaël Leblois, Pierre-André Crochet

## Abstract

**Aim:** Discontinuity in the distribution of genetic diversity (often based on mtDNA) is usually interpreted as evidence for phylogeographic breaks, underlying vicariant units. However, a misleading signal of phylogeographic break can arise in the absence of barrier to gene flow, under mechanisms of isolation by distance (IBD). How and under which conditions phylogeographic breaks can be reliably differentiated from populations evolving under IBD remain unclear. Here, we use multi-locus sequence data from a widely distributed lizard species to address these questions in an empirical setting.

**Location:** Morocco

**Taxon:** Spiny-footed lizard (*Acanthodactylus erythrurus*), Squamata: Lacertidae

**Methods:** Using 325 samples from 40 localities, we identified genetic discontinuities within *A. erythrurus* based on a mitochondrial fragment and nine nuclear markers independently. Using the nuclear markers, we then applied linear regression models to investigate whether genetic divergence could be explained by geographical distances alone, or barriers to gene flow (real phylogeographic breaks).

**Result:** *A. erythrurus* is characterized by an important mitochondrial diversity, with 11 strongly supported phylogeographic lineages with a crown age of 6 Mya. Nuclear markers, however, yielded weak phylogenetic support for these lineages. Using clustering methods based on genotypes at nine nuclear loci, we identified phylogeographic clusters that were partly discordant with the mtDNA lineages. Tests of IBD delimited at least four groups of populations separated by barriers to gene flow, but unambiguous separation of vicariance from IBD remained challenging in several cases.

**Main conclusions:** The genetic diversity of *A. erythrurus* originates from a mix of IBD and vicariance, which were difficult to distinguish, and resulted in similar levels of mitochondrial differentiation. These results highlight that phylogeographic breaks inferred from mitochondrial data should be further investigated using multi-locus data and explicit testing to rule out alternative processes generating discontinuities in mitochondrial diversity, including IBD. We identified four groups of populations within *A. erythrurus*, separated by barriers to gene flow, but even using nine independent nuclear makers the power of our approach was limited, and further investigation using genome-wide data will be required to resolve the phylogeographic history of this species.

## Introduction

One of the main goals of phylogeography is to identify discontinuities in the distribution of genetic diversity across conspecific populations – phylogeographic breaks – and investigate the biotic and abiotic factors underlying them (Kumar & Kumar, 2018). Typically, phylogeographic breaks are interpreted as evidence for the existence of evolutionary lineages that diverged through vicariance events (Kuo & Avise, 2005). As such, they have been widely used to investigate biogeography and evolutionary biology questions, including the impact of geological events and landscape features on populations connectivity (e.g., Leaché, Oaks, Ofori-Boateng & Fujita 2020), and their respective roles in promoting divergence and speciation (Huang, 2020). Furthermore, phylogeographic breaks are of interest for other fields of study, such as taxonomy and conservation biology (Braby, Eastwood & Murray 2012; Shaffer et al., 2015). However, correct identification of phylogeographic breaks remains a non-trivial problem (Audzijonyte & Vrijenhoek, 2010).

Historically, phylogeographic studies have heavily relied on uniparentally inherited markers, such as mitochondrial DNA (mtDNA), to infer patterns of genetic diversity (Avise et al., 1987). However, while being characterized by a number of advantages (e.g., high mutation rate, lower effective population size, established standardized sequencing protocols), these markers often yield misleading signals of phylogeographic breaks (or the absence thereof) due to a number of processes (reviewed in Toews & Brelsford, 2012). Among them, different selective pressures (Irwin, 2012), introgression (e.g., Zieliński et al., 2013) and sex-biased dispersal (Palumbi & Baker, 1994; Peters, Bolender & Pearce, 2012; Dávalos & Russell, 2014) are well documented, while incomplete lineage sorting is not expected to generate predictable, biogeographically discordant patterns (Funk & Omland, 2003; Toews & Brelsford, 2012). Conversely, breaks in mitochondrial genetic diversity can also appear in continuous populations solely as a result of isolation by distance (IBD; Irwin, 2002), especially in the presence of sampling gaps (Audzijonyte & Vrijenhoek, 2010), and mtDNA is often poorly suited to detect IBD patterns (Teske et al., 2018). Additionally, mtDNA gene trees often differ from the population tree, mainly due to introgression and incomplete lineage sorting (ILS), hence phylogeographic breaks identified from mtDNA might not reflect actual populations splits (e.g., Dufresnes, Nicieza et al., 2020; Velo-Antón, Martínez-Freiría, Pereira, Crochet & Brito 2018).

The limitations of mitochondrial phylogeography are usually overcome by considering multiple unlinked nuclear loci, which enable more accurate inferences (Brito & Edwards, 2009), particularly in the framework of the coalescent theory (Knowles, 2009). This approach reduces the likelihood of detecting spurious phylogeographic breaks, as independent loci are not expected to support concordant breaks in the absence of barriers to gene flow (Avise & Ball, 1990; Kuo & Avise, 2005). However, populations evolving along a single geographical gradient (i.e., under IBD) might still be apparently divided into distinct lineages, for example in the presence of large sampling gaps (Schwartz & McKelvey, 2009; Tolley et al., 2022; Wiemers & Fiedler, 2007). In that context, it appears crucial to test explicitly whether apparent phylogeographic breaks (e.g., reciprocally monophyletic groups in phylogenetic trees) delimit vicariant units separated by barriers to gene flow or have been generated by alternative processes such as IBD. A commonly used approach is to study the correlation between genetic and geographical distances among populations or individuals in a statistical framework (e.g., Ramachandran et al., 2005), to determine whether the null hypothesis that all populations differentiate along the same IBD gradient can be rejected, which would indicate the presence of a barrier to gene flow (e.g., Binks et al., 2019; Hausdorf & Hennig, 2020; Goudarzi et al., 2019). Other popular approaches include the estimation of migration rates over geographical surfaces, with low migration rates highlighting barriers to gene flow (Petkova, Novembre & Stephens, 2016). However, in practice, the implementation of such tests is often limited in phylogeographic studies by sparse sampling schemes and the low number of genetic markers available. Here, we investigate the extent to which vicariant units can safely be identified from populations evolving along IBD gradients, using multi-locus data of a genetically diverse North African lizard.

Compared to other regions of the Mediterranean Basin biodiversity hotspot, fewer phylogeographic studies have focused on North African communities (Beddek et al., 2018; Husemann, Schmitt, Zachos, Ulrich & Habel, 2014). Among these, many studies have revealed important intra-specific genetic diversity in various taxa (reviewed in Husemann et al., 2014), in relation with the range of landscapes and bioclimatic regions and the complex biogeographic history that characterize this area. Consequently, important gaps still hamper our knowledge of the taxonomic diversity of North African taxa and the evolutionary processes which created it (Beddek et al., 2018; Brito et al., 2014; Ficetola, Bonardi, Sindaco & Padoa-Schioppa, 2013).

Phylogeographic studies of North African taxa are needed to fill the aforementioned knowledge gaps, by documenting the extent and geographic distribution of intra-specific genetic diversity, and identifying unrecognized species-level lineages. Secondly, biodiversity hotspots such as North Africa are prime areas to study the ecological and evolutionary processes shaping the distribution of genetic diversity and underlying lineages diversification at a global scale (e.g., Carnaval, Hickerson, Haddad, Rodrigues & Moritz, 2009; Demos, Peterhans, Agwanda & Hickerson, 2014; Durrant, Barrett, Edgar, Coleman & Burridge, 2015). In that context, regionally distributed (as opposed to narrow-ranged) taxa are particularly valuable models, as they are likely to inform on a larger variety of processes.

In this study, we focus on the Red-tailed Spiny-footed lizard *Acanthodactylus erythrurus* (Squamata: Lacertidae), a species widely distributed across the Maghreb and the Iberian Peninsula. This species is characterized by pronounced morphological variations that led to the recognition of several morphological subspecies or species (Salvador, 1982; Bons & Geniez, 1995), which inhabit distinct habitat types (from coastal sand dunes to high altitude xerophytic vegetation through open forests and semi-arid steppes). Several molecular studies (Beddek et al., 2018; Fonseca, Brito, Paulo, Carretero & Harris, 2009; Miralles et al., 2020) have revealed an important and previously unsuspected intraspecific genetic diversity, leading to the description of two new species, and emphasizing the need for a more in-depth taxonomic revision of this group. Notably, these studies highlight important disagreements between the accepted taxonomy and molecular clades. One of the main questions raised by these findings is whether the morphological and ecological diversification of this group has been produced by vicariance and lineage diversification, or by local adaptation in the face of historical gene flow. However, most studies (except Miralles et al., 2020) have relied only on mtDNA, offering an incomplete picture of the historical processes underlying *A. erythrurus*’ genetic diversity. Here, we complement the multi-locus data produced by Miralles et al. (2020) with new sequences, focusing on Moroccan populations with a comprehensive sampling, to clarify whether multiple vicariant units explain the genetic diversity in the *A. erythrurus* complex (referred to as *A. erythrurus* here).

The goals of this study are 1) to identify candidate phylogeographic breaks among Moroccan populations of *A. erythrurus* using both mtDNA and nuclear DNA (nDNA) markers; 2) to determine whether these candidate breaks delimit vicariant units or could result from IBD, and 3) integrate these processes into a phylogeographic scenario to explain *A. erythrurus*’ present genetic diversity. Across the range of *A. erythrurus,* we decided to focus on Moroccan populations to mitigate the effect of sampling gaps, as we only had sparse sampling in other countries. We employed a two-step approach to untangle the processes at play: first, we analyzed mitochondrial and nuclear (nine gene sequence fragments) data independently to identify phylogeographic lineages. In both cases, phylogeographic lineages were defined as the smallest genetically homogeneous geographical units. Secondly, we used the nuclear dataset to test whether the described lineages corresponded to vicariant units or could result from IBD alone. Through this approach, we aim at providing a more comprehensive picture of the phylogeographic history of Moroccan *A. erythrurus*, but also empirically assess our ability to identify vicariant units using multi-locus data in a genetically variable species.

## Material & Methods

### 1. Sampling and DNA sequencing

We used a multi-locus dataset of 325 samples from the Moroccan populations of *A. erythrurus.* Among these, 128 samples have been previously used by Miralles et al. (2020) in their revision of the whole *A. erythrurus* complex using samples from its whole range (Algeria, Morocco, Portugal, Spain, Tunisia). The remaining 197 have never been analyzed in any publication so far. In the present study, we focus on the Moroccan populations of *A. erythrurus* because they offer the most complete sampling. These 325 samples represent the Moroccan localities of the “Ibero-Moroccan” (IM) clade defined by Miralles et al. (2020), but with significantly more samples per locality. The other Moroccan populations historically assigned to *A. erythrurus* (Western and Eastern High Atlas clades of Miralles et al., 2020) represent distinct species described as such by Miralles et al. (2020). They are isolated from the IM clade by strong barriers to gene flow and were thus not considered here. Ten additional samples from other clades and species were also included to be used as outgroups in phylogenetic analyses, as described below. Details on the localities and number of samples for the present study are given in Figure 1 and Table S1.

**Figure 1.**
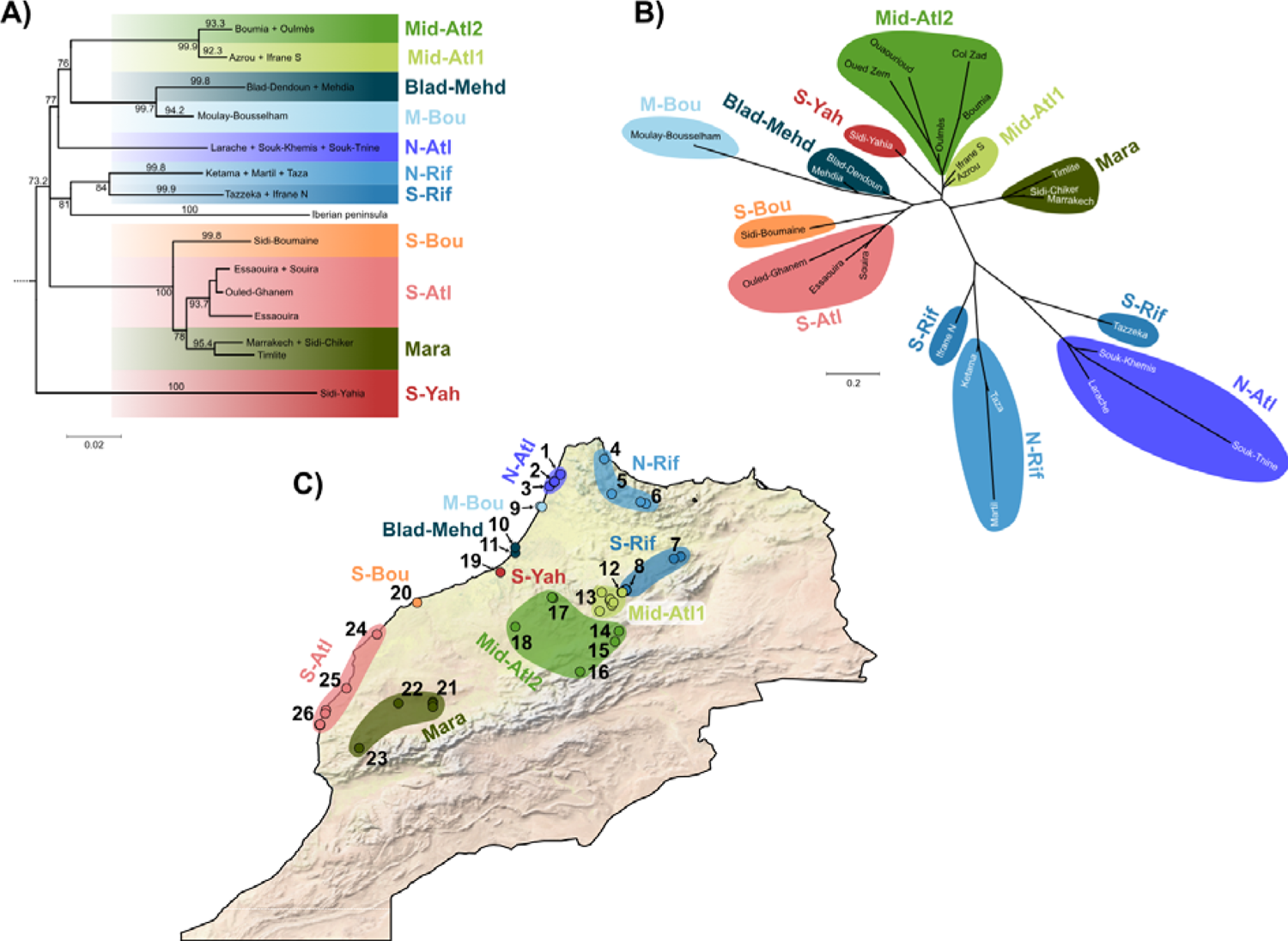
A) Simplified phylogenetic tree obtained from the maximum likelihood analysis of 776 bp of the ND4 mitochondrial fragment. The outgroups used for rooting (10 samples of closely related A. erythrurus lineages and five other Acanthodactylus species) were removed from the figure and terminal groups were pruned for improved graphical resolution (for full tree, see *Figure S1*).

All samples have been sequenced for DNA fragments from one mitochondrial gene (776 bp of the NADH-dehydrogenase subunit 4 [ND4] gene) and nine nuclear protein-coding genes: Recombination Activating Gene 2 (RAG2, 321 pb); Oocyte maturation factor (C-mos, 369 pb), Ornithine Decarboxylase (OD, 396 pb), Phosphogluconase Dehydrogenase intron 7 (PgD7, 420 pb), Melanocortin Receptor 1 (MC1R, 630 pb), Pyruvate Kinase Muscle (PKM, 351 pb), Nuclear transcription factor Y gamma (NFYC, 666 pb), Acetylcholinergic receptor M4 (ACM4, 390 pb) and RNA fingerprint protein 35 (R35, 588 pb). Detailed sampling and laboratory protocols are given in Miralles et al. (2020). All sequences were submitted to Genbank, and individual accession numbers are available in Appendix 1.

Finally, for the ingroup samples, individual genotypes were inferred at each nuclear locus using the software PHASE (Stephens, Smith & Donnelly, 2001) implemented in DNAsp v. 5 (Librado & Rozas, 2009). The samples were grouped based on their mitochondrial lineages (cf. Results) to account for the assumption that all samples are drawn from a single population at Hardy-Weinberg equilibrium (Stephens et al., 2001). Such approach could be problematic if individuals from separated populations shared closely related mtDNA haplotypes (e.g., in the case of mitochondrial introgression between two sampled populations). However, preliminary analyses confirmed that samples sharing similar mtDNA haplotypes were always closely related in the nDNA data. PHASE was run on each group separately, with 5,000 sampled steps and 10,000 burnin-steps in the MCMC. Uncertain phases (i.e., Posterior Probability < 0.90) were treated as unresolved sites using PHASE’s default settings.

### 2. Mitochondrial phylogeography

As a first step to identify the lineages used as basic units for the subsequent analyses, we conducted phylogenetic inference based on 776 bp of the ND4 mitochondrial gene. In addition to all our Moroccan samples, two samples from Portugal and Spain were also included to allow estimation of the divergence times based on the opening of the Gibraltar straight (more details below). Finally, the tree was rooted by including representatives of each of the following lineages (for the lineages nomenclature, we follow Miralles et al., 2020): *A. savignyi* (N=1), *A. margaritae* (N=1), *A. lacrymae* (N=2), the AT lineage from eastern Algeria and Tunisia (including *A. blanci,* N=2), *A. montanus* (N=2) and the CA lineage from central Algeria (N=2). In total, 331 samples were included in the final alignment (ND4 sequencing failed for four ingroup samples).

Maximum-likelihood (ML) phylogenetic inference was performed using IQTREE v. 1.6.8 (Chernomor, von Haeseler & Minh, 2016; Nguyen, Schmidt, von Haeseler & Minh, 2015). The best substitution model was selected using ModelFinder (Kalyaanamoorthy, Minh, Wong, von Haeseler & Jermiin, 2017) as implemented in IQTREE, and the branches support was assessed using the SH-like approximate likelihood test (aLRT) with 5,000 pseudoreplicates. Based on this tree, we defined phylogeographic lineages corresponding to the smallest geographical units supported as monophyletic groups (in practice, with an aLRT support > 90). Finally, we estimated divergence times through two approaches. First, we calibrated the divergence between the Iberian populations and their closest Moroccan relatives based on the opening of the Strait of Gibraltar (between 5.3 and 5.5 Million years ago [Mya]). However, as more recent colonization events across the Strait of Gibraltar have been found in reptiles and amphibians (e.g., Carranza, Harris, Arnold & Gonzalez de la Vega, 2006; Gutiérrez-Rodríguez, Márcia Barbosa & Martínez-Solano, 2017), we also calibrated the tree based on a fixed mutation rate for the ND4 gene. Details on the procedure can be found in Supplementary materials S1.

### 3. Nuclear phylogeography

Preliminary analyses revealed that the phylogeographic signal in the nuclear data was much weaker compared to that of the mitochondrial dataset, with much higher levels of allele sharing among populations and lower nodes support in phylogenies. Our initial approach to infer genetic clusters and admixture among them was to use the software STRUCTURE (Pritchard, Stephens & Donnelly, 2000). However, we failed to reach proper convergence of the MCMC chains and to identify the optimal number of clusters (K). Furthermore, for a single K value, independent runs yielded very different, yet equally likely, clustering solutions (results not included). These solutions suggested clearly separated groups (i.e., high Q values), but their composition varied across runs, and did not highlight a clear geographic signal. Likewise, phylogenetic reconstructions of the concatenated data failed to converge toward a consistent topology. In that context, it was difficult to apply a strict criterion to delimit phylogeographic lineages from the nuclear dataset. Thus, we relied on congruent results from several analyses, especially multivariate analyses of the genotypes at the nine nuclear loci, as described below.

### 3.1 Phylogenetic analyses

We first performed ML phylogenetic inference using the concatenation of the nine nuclear loci. Best substitution models and partition schemes were selected on a per-locus partition, and the analyses were run in IQTREE v. 1.6.8 with the same settings as for the mitochondrial analyses. Five independent ML searches were performed.

In addition, we also performed Neighbour-Joining (NJ) inference based on between-populations (i.e., sampling localities) pairwise distance matrices for the ingroup samples only. This analysis was repeated twice, firstly using average p-distances calculated from the concatenation of the nine markers, secondly using Nei’s distances computed from the genotypes at the nine loci. In the first case, both distances computation and tree inference were performed in MEGA 11 (Tamura, Stecher & Kumar, 2021); in the second case Nei’s distances were calculated using adegenet v. 2.1.5 (Jombart, 2008) and NJ inference performed with ape v. 5.6 (Paradis & Schliep, 2019).

Because our attempts to infer a phylogenetic tree from the nuclear markers yielded poorly supported and variable topologies, in which the position of the Iberian samples is different from that of the mitochondrial tree (see Results and Figure S5), we did not attempt to calibrate the nuclear phylogeny. Furthermore, we spent a substantial amount of time trying to get a species tree from the nine nDNA markers using *BEAST (Heled & Drummond, 2009) but did not manage to reach proper convergence and obtained variable topologies across runs.

### 3.2 Multivariate analyses

As an alternative to phylogenetic analyses, we used Multiple Correspondence Analyses (MCA) to detect phylogeographic breaks based on the nuclear dataset. MCAs were calculated from the genotypes at the 9 nuclear loci, using the software Genetix (Belkhir, Korsa, Chikhi, Raufaste & Bonhomme, 2004). We used an iterative approach to define phylogeographic lineages: the analysis was first run on a dataset including all Moroccan populations (same sampling as for the phylogenetic analyses but without outgroups and Iberian samples); the results were visually inspected and groups of populations that were clearly differentiated from the rest were identified as phylogeographic lineages. These populations were then removed from the dataset, and the analysis was run again on the resulting subset. This process was repeated until no clear structure could be identified in the results (in total, five iterations were needed, see Results).

### 4. Tests of Isolation by Distance and vicariance

Population structure analyses based on both mtDNA and nDNA data supported several genetic groups in Moroccan *A. erythrurus.* While the structure inferred from both types of markers was mostly concordant, there were a few notable differences. For further analyses, we considered the nDNA genetic groups (summarized in the following, cf. Table 1 and Figure 2 for more details), which minimized the number of clusters represented by a single locality. Indeed, the employed statistical test relies on calculating a regression on within-group geographic distances, which is not possible for single localities. More precisely, we defined four inland groups: the Rif Mountains (Rif, including N-Rif and S-Rif), the Middle-Atlas Mountains (Mid-Atl, including Mid-Atl1 and Mid-Atl2), the surroundings of Marrakech (Mara), and the population of Sidi-Yahia (S-Yah); as well as six groups distributed along the Atlantic coast, most of which consist in single localities, except for the northernmost (N-Atl) and southernmost (S-Atl) groups. To determine whether this pattern of genetic diversity reflects true phylogeographic breaks, we investigated whether these genetic groups differentiate along similar geographical gradients (i.e., IBD gradients). The populations were divided into two groups, analyzed separately for practical purposes: 1) the inland lineages and 2) the lineages distributed along the Atlantic coast. All scripts and data files used to run these analyses are available at https://github.com/rancilhac/IBD_vs_Vicariance (http://doi.org/10.5281/zenodo.7142595).

**Figure 2.**
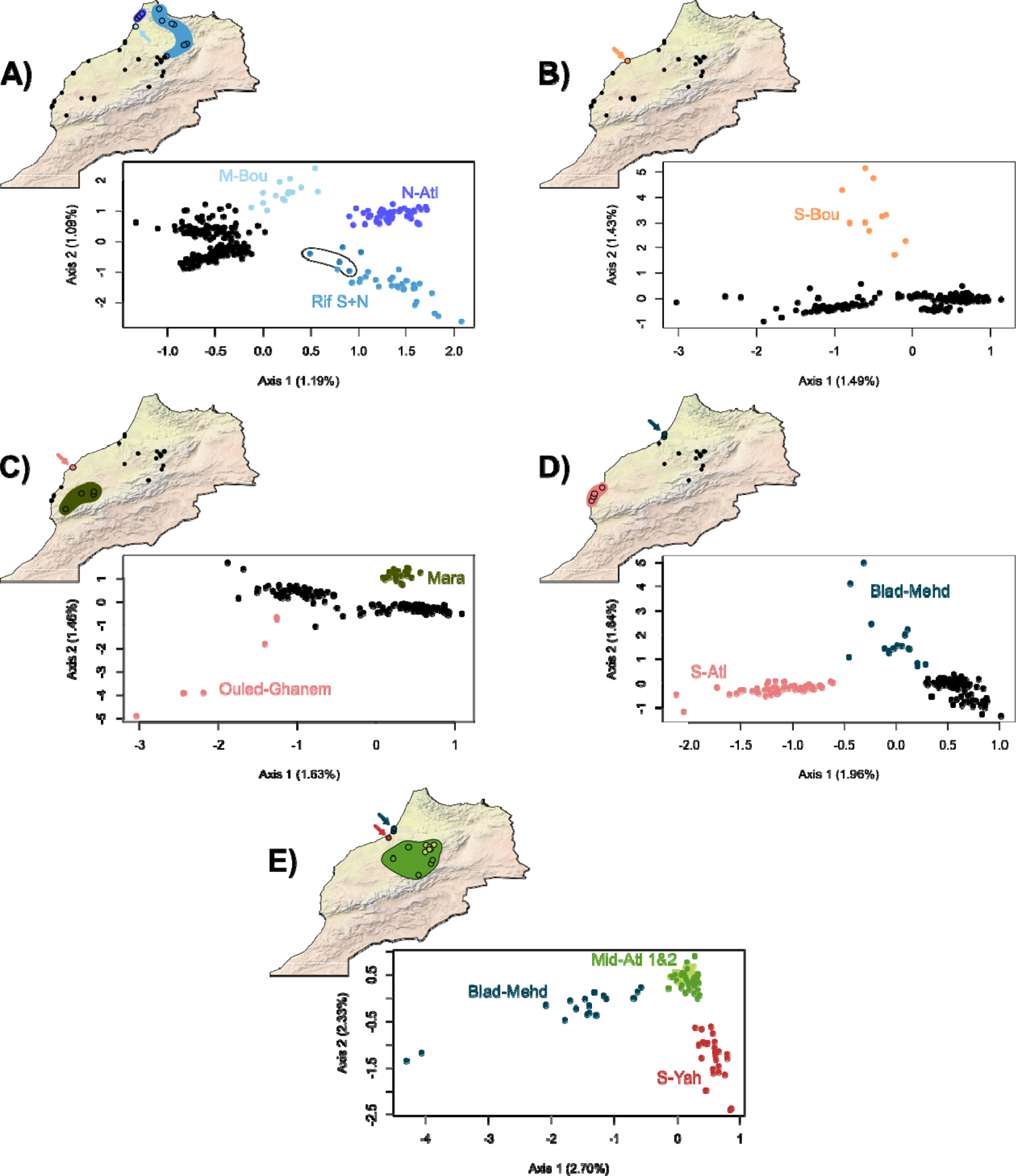
Results of Multiple Correspondence Analyses calculated from the genotypes at the nine nuclear genes. The analysis was repeated five times iteratively with subsets of populations to detect finer phylogeographic structure. Colors and names correspond to the mitochondrial lineages. In A, the black ellipse highlights three samples from the contact zone between Rif and Mid-Atl. Blad-Mehd is highlighted twice in D and E to illustrate its intermediate position between the Southern Atlantic and Middle-Atlas populations.

**Table 1.** Summary of the phylogeographic lineages delimited using 776 bp of the ND4 mitochondrial gene, and genotypes at nine nuclear loci. Numbers between brackets in the fourth column refer to the localities numbering on *Figure 1C* and *Table S1*. The full sample list for each locality is given in *Appendix 1*.

| Mitochondrial lineages | Nuclear lineages | Geographic range | Localities |
| --- | --- | --- | --- |
| N-Atl | N-Atl | Northernmost Atlantic coast | Larache, Souk-Khemis, Souk-Tnine [1-3] |
| N-Rif | Rif | Northern part of the Rif mountains | Ketama, Martil, Bab-Taza [4-6] |
| S-Rif |  | Northern parts of Middle-Atlas | Tazzeka, Ifrane North [7-8] |
| M-Bou | M-Bou | Single locality on the North of the Atlantic coast south of Larache | Moulay-Bousselam [9] |
| Blad-Mehd | Blad-Mehd | Two neighbour localities on the Atlantic coast just north of Rabat | Blad dendoun, Mehdia [10-11] |
| Mid-Atl1 | Mid-Atl | North-eastern parts of the middle-Atlas | Ifrane South, Azrou area [12-13] |
| Mid-Atl2 |  | Southern and western parts of the middle-Atlas | Oulmès area, Boumia, Col Zad, Ouaouriod, Oued Zem [14-18] |
| S-Yah | S-Yah | Single locality just south of Rabat | Sidi-Yahia [19] |
| S-Bou | S-Bou | Single locality on the Atlantic coast south of Casablanca | Sidi-Boumaine [20] |
| Mara | Mara | Populations surrounding Marrakech, in southern Morocco | Marakech area, Sidi Chiker, Timlite [21-23] |
| S-Atl | O-Gha | Single locality on the Atlantic coast north of Essaouira | Ouled Ghanem [24] |
|  | S-Atl | Populations of the southernmost Atlantic coast | Essaouira area, Souira [25-26] |

### 4.1 Differentiation between inland lineages

To assess whether the Rif, Mid-Atl and Mara groups are vicariant units, we analyzed the correlation between genetic and geographical distances in pairs of geographically adjacent lineages, using the statistical framework of Hausdorf & Hennig (2020). This approach relies on the null hypothesis that the two considered groups of population differentiate along the same IBD gradient. Testing this relies on a two-steps approach: firstly, we tested that the within-groups distances of both lineages followed similar regressions (hypothesis *H01* of Hausdorf & Hennig, 2020). In case this was verified, we then tested whether regressions fitted on between-groups distances only and all within-group distances are similar (hypothesis *H02* of Hausdorf & Hennig, 2020). A rejection of that hypothesis (i.e. within-groups and between-groups distances do not fit the same regression line) would support a barrier to gene flow between the tested groups. However, this test is not appropriate if *H01* is not verified. In such case, we tested whether between-groups distances follow the same regression as the within-group distances of one of the two lineages (hypothesis *H03* of Hausdorf & Hennig 2020). A verification of that hypothesis is incompatible with a barrier to gene flow. When testing *H03*, we performed two tests, *H031* and *H032,* in which the within-group distances used for the comparison are respectively those of group 1 and group 2. A failure to reject both *H031* and *H032* would indicate inconclusive results deserving further investigation (Hausdorf & Hennig, 2020). In all cases, the significance of the difference between regression lines is assessed using a non-parametric jackknife test, as described in Hausdorf & Hennig (2020). We used this approach to test the following combinations of lineages, corresponding to the inland lineages and adjacent coastal lineages: Rif + Atl-N, Rif + Mid-Atl, Mid-Atl + Mara, Mid-Atl + (S-Atl, O-Gha, S-Bou), Mara + S-Atl.

The case of S-Yah is more complicated to address, because this lineage is represented by a single locality, thus the tests described in the previous section are not applicable. Considering the results of previous analyses (cf. Results, Figure 2), we hypothesize that S-Yah is closely related to the Mid-Atl group. To test whether S-Yah differentiate within the same IBD gradient as Mid-Atl, we tested whether regressions within (Mid-Atl + S-Yah) and within Mid-Atl alone were similar. If this hypothesis is verified, we interpret it as an indication for an absence of barrier to gene flow between Mid-Atl and S-Yah.

All regression tests were performed using the prabclus R package (Hausdorf & Hennig, 2020) based on pairwise distances between individuals. In the specific case of S-Yah, the *regdistdiff* function had to be adapted in a custom R script to match our needs. Geographic distance matrices were calculated from coordinates using prabclus, and log transformed. Genetic distance matrices were computed from the genotype data at the nine nuclear loci using two metrics: the *â* statistic (an equivalent of *Fst*/(1-*Fst*) calculated between individuals), calculated in Genepop (Rousset, 2008), and the shared allele distance (*s*), calculated in prabclus. Each analysis was repeated twice, using both measures of genetic distance. We decided not to use mitochondrial distances because empirical evidence suggest that mitochondrial markers are not suited to detect IBD gradients (Audzijonyte & Vrijenhoek, 2010; Teske et al., 2018). Furthermore, male-biased dispersal has been identified in most Squamates studied to date (Ferchaud et al. 2015), meaning that maternally inherited markers might not accurately represent dispersal patterns in *A. erythrurus*.

### 4.2 Isolation by distance along the Atlantic coast

Because several of the genetic groups along the Atlantic coast are represented by single localities, testing for the presence of barriers to gene flow between each group pair is not straightforward, as explained in the case of S-Yah (above). Thus, rather than testing each putative phylogeographic breaks, we explored two hypotheses. First, considering that N-Atl is very distinct from other coastal populations on multivariate analyses and population trees (where it appears to be more related to the Rif, cf. Results), we hypothesize that there is a barrier to gene flow between N-Atl and the population adjacent to the south, M-Bou. Testing this hypothesis is relatively straightforward as the framework of Hausdorf & Hennig 2020 applies here, by testing (S-Atl, O-Gha, S-Bou, Blad-Mehd, M-Bou) and N-Atl. However, it is also possible that there are more barriers to gene flow along the coast, or none. To gain more insight into this, we first calculated a regression between genetic and logarithm of geographical distances including all coastal lineages (to the exclusion of N-Atl in case the previous test showed that it does not evolve along the same IBD gradient as the southernmost populations). We then successively removed lineages and re-calculated the regression. In case removing populations did not significantly change the regression, we interpret this as an indication that all coastal lineages differentiate along a single IBD gradient. All tests were performed in the same set up as detailed in the previous section.

## Results

### 1. Mitochondrial DNA supports deep phylogeographic lineages in Moroccan Acanthodactylus erythrurus

Using 776 bp of the ND4 mitochondrial gene, we recovered a relatively well supported phylogenetic tree (Figure 1, Figure S1), fully consistent with the analyses of the same data by Miralles et al. (2020). Particularly, the monophyly of the “Ibero-Moroccan” (IM) clade is fully supported (aLRT = 100), and this group is further divided into 11 phylogeographic lineages with aLRT support >90 (Figure 1, Table 1), namely (from north to south): 1) the northern Atlantic coast (N-Atl, from Souk-Tnine to Larache); 2) the northern part of the Rif mountains (N-Rif, from Martil to Ketama); 3) the southern part of the Rif (S-Rif, Tazzeka and the North of Ifrane); 4) the coastal population of Moulay-Bousselham (M-Bou); 5) the coastal populations of Blad-Dendoun and Mehdia (Blad-Mehd); 6) the inner-most parts of the Middle-Atlas mountains (Mid-Atl1, around Azrou and Ifrane); 7) the rest of the Middle-Atlas mountains (Mid-Atl2, from Oulmès to Ouaourioud and Boumia); 8) the population of Sidi-Yahia (S-Yah); 9) the coastal population of Sidi-Boumaine (S-Bou); 10) the populations surrounding Marrakech (Mara; including Sidi-Chiker and Timlite) and 11) the Southern Atlantic coast (S-Atl; from Ouled-Ghanem south to Essaouira). All lineages are geographically isolated by sampling gaps, except for S-Rif and Mid-Atl1, which come into contact north of Ifrane (Middle-Atlas). Of the three samples from this locality, two cluster with the former and one with the latter (see Doniol-Valcroze et al. 2022 and Figure S1; the correspondence between samples and localities is given in Appendix 1).

The 11 lineages cluster into three main clades: i) Mara, S-Atl, S-Bou; ii) Rif N, Rif S (as well as the Iberian samples) and iii) N-Atl, M-Bou, Blad-Mehd, Mid-Atl1, Mid-Atl2. The aLRT support for these clades ranges from 77 to 100, but the relationships among them remain unresolved. Finally, S-Yah harbors a remarkably divergent mitochondrial lineage which is basal to all other Moroccan mitochondrial clades of A. erythrurus. Numbers at the nodes show aLRT support Numbers on terminal branches show the aLRT support of collapsed terminal clades. Color boxes show the 11 identified phylogeographic lineages. B) Neighbour-Joining population tree inferred from the pairwise Nei’s distances between populations, calculated from the genotypes at nine nuclear loci, with colors referring to the mitochondrial lineages. C) Map of Morocco showing the sampled localities, with colors and texts showing the mitochondrial lineages they belong to. Localities are numbered as in Table 1, with coordinates and sample sizes given in Table S1.

The time-tree analyses based on the end of the Messinian crisis (Figure S2) inferred an origin of the “IM” clade at 5.99 Mya (95% credibility interval [5.36 – 7.07]). Inferences with a fixed mutation rate (Figures S3 & S4) yielded slightly older dates (6.05 and 7.12 Mya based on the mutation rate of *Podarcis* and *Gallotia* respectively), but the 95% confidence intervals were much wider. Overall, the three calibration approaches gave similar dates. However, these results should be taken with care since the analyses failed to reach proper convergence (see Supplementary materials S1 for more details).

### 2. Multi-locus nuclear phylogeography

The ML phylogenetic analysis of the concatenation of the nine nuclear markers yielded a much less resolved topology compared to that of the mtDNA data (Figure S5). While the monophyly of the IM clade is strongly supported (aLRT = 99), most of the mitochondrial lineages are mixed and do not form reciprocally monophyletic groups. However, the following mitochondrial lineages are also supported as monophyletic in the nuclear phylogeny: Mara (aLRT = 97); S-Bou (aLRT = 81); and a clade composed of M-Bou and N-Atl (aLRT = 90) within which the former is paraphyletic but the latter is monophyletic (aLRT = 91). Finally, a clade comprising all N-Rif and S-Rif samples to the exclusion of one from the contact zone with Mid-Atl1 is relatively well supported (aLRT = 75), but also includes one sample from Mid-Atl1.

Multiple Correspondence Analyses (MCA) of the genotypes at the nine nuclear loci recovered a phylogeographic structure partially concordant with that of the mitochondrial data (Figure 2). With this approach, we could identify 10 phylogeographic lineages (from north to south, the names refer to those used for the mitochondrial lineages, see also Table 1): 1) N-Atl; 2) the Rif mountains (Rif, mitochondrial lineages N-Rif & S-Rif; including all three samples from IFRA-N); 3) M-Bou; 4) Blad-Mehd; 5) all the Middle Atlas populations (Mid-Atl, mitochondrial lineages Mid-Atl1 & Mid-Atl2); 6) S-Yah; 7) S-Bou; 8) Mara; 9) the coastal locality of Ouled-Ghanem (O-Gha, grouped within the S-Atl lineage in the mitochondrial data); and 10) S-Atl (to the exception of O-Gha). Overall, these groups are mostly overlapping with the mitochondrial lineages, but are less straightforward to delimit due to the weak phylogenetic resolution of the nuclear markers. The locality O-Gha appears as an exception, being clustered with the other southern Atlantic populations in the mitochondrial data but quite divergent in the nuclear data (Figure 2C). Interestingly, S-Yah appears here closely related to Mid-Atl populations, from which it could be distinguished only at the finer scale (Figure 2E), contrasting with its high mitochondrial divergence.

Population NJ trees inferred from both p-distances (Figure S4) and Nei distances (Figure 1B) supported a structure similar to that of the MCA. The Rif, N-Atl and Mara groups are separated from the others by relatively long branches (although Mara is closer to the other populations than to Rif and N-Atl), while populations from the southern Atlantic coast, Mid-Atl and S-Yah appear closely related. Both genetic distances did not yield completely concordant topologies, especially regarding the position of M-Bou, which is alternatively placed close to N-Atl or S-Atl. On both trees, Blad-Mehd has an intermediate position, suggesting an effect of admixture between several genetic groups.

### 3. Isolation by Distance tests

IBD tests for pairs of inland lineages based on both *â* and *s* genetic distances were overall congruent (Tables 2 & 3). Figures 3 & 4 display the regression lines based on *â*, while those based on *s* are provided in Figures S7 & S8.

**Figure 3.**
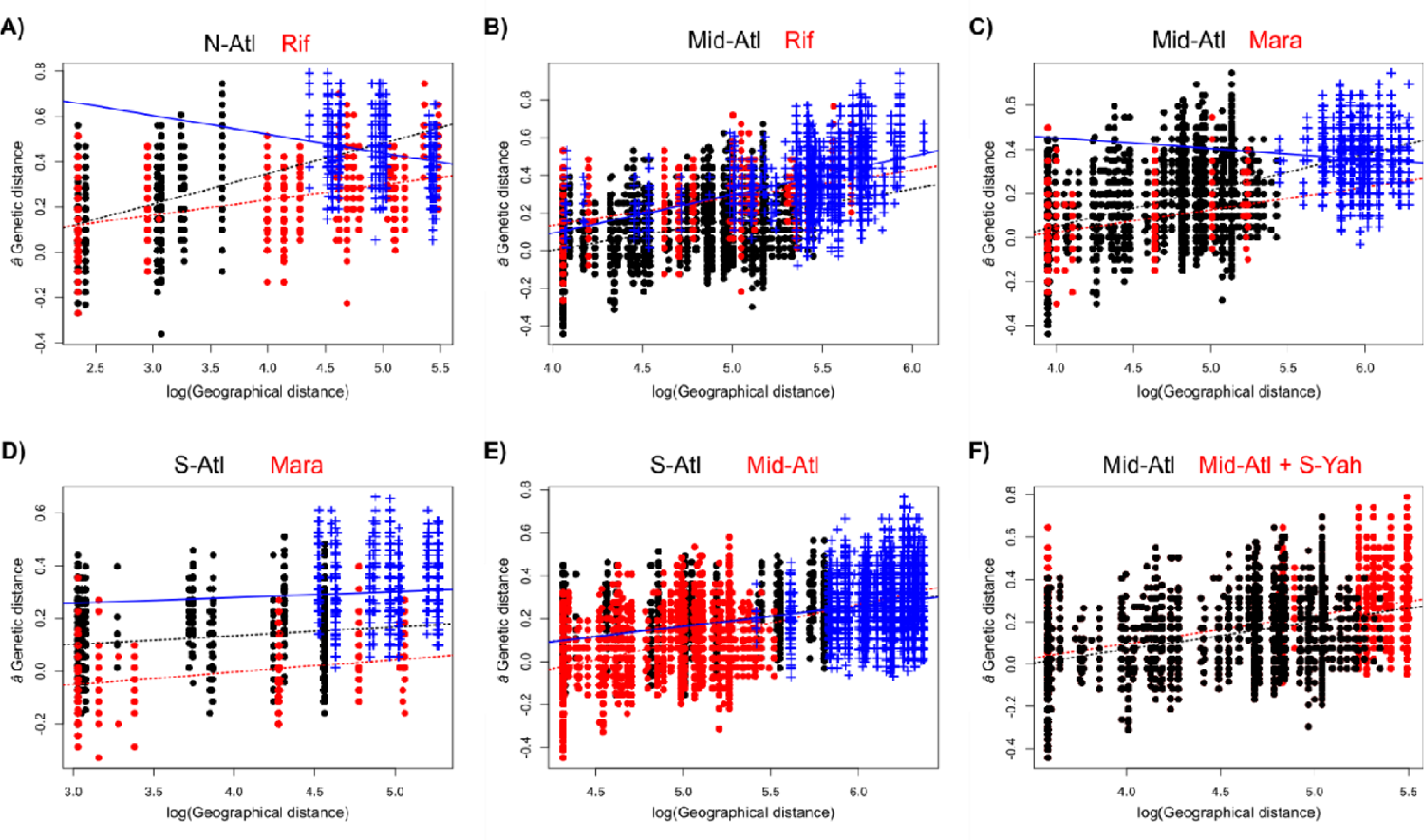
Linear regressions between genetic (â) and geographical distances in pairs of geographically adjacent inland lineages. Black and red dots show within-group distances for both lineages, and corresponding dashed lines represent linear regression fitted to them. Blue crosses and solid lines show between-group distances. Between-group distances are not represented on the last graph as the test was adapted to account for zero-only geographical distances in S-Yah.

**Table 2.** Results of isolation by distance tests (p-values) based on the comparisons of between-groups and within-groups distances. H01, H02, H031 and H032 refer to the different hypotheses tested, as explained in the text. â = â genetic distance; s = shared allele distance.

|  |  | *H01* |  | *H02* |  | *H031* |  | *H032* |  |
| --- | --- | --- | --- | --- | --- | --- | --- | --- | --- |
| Group 1 | Group 2 | $\hat{a}$ | *s* | $\hat{a}$ | *s* | $\hat{a}$ | *s* | $\hat{a}$ | *s* |
| *Inland lineages* |  |  |  |  |  |  |  |  |  |
| N-Atl | Rif | 0.35 | 0.40 | 5.69e-6 | 6.58e-6 | - | - | - | - |
| Rif | Mid-Atl | 2.96e-4 | 0.08 | - | 3.93e-7 | 2.17e-6 | - | 0.19 | - |
| Mid-Atl | Mara | 0.07 | 3.94e-5 | 0.55 | - | - | 0.91 | - | 0.025 |
| Mara | S-Atl | 1.25e-5 | 3.6e-13 | - | - | 4.14e-8 | 0.006 | 6.20e-8 | 6.48e-14 |
| S-Atl | Mid-Atl | 0.0005 | 0.0006 | - | - | 0.50 | 0.49 | 0.73 | 0.74 |
| *Coastal lineages* |  |  |  |  |  |  |  |  |  |
| N-Atl | (M-Bou + Bad-Mehd + S-Bou + O-Gha + S-Atl) | 6.05e-20 | 1.33e-15 | - | - | 7.94e-13 | 3.02e-5 | 1.00 | 1.00 |

**Table 3.**
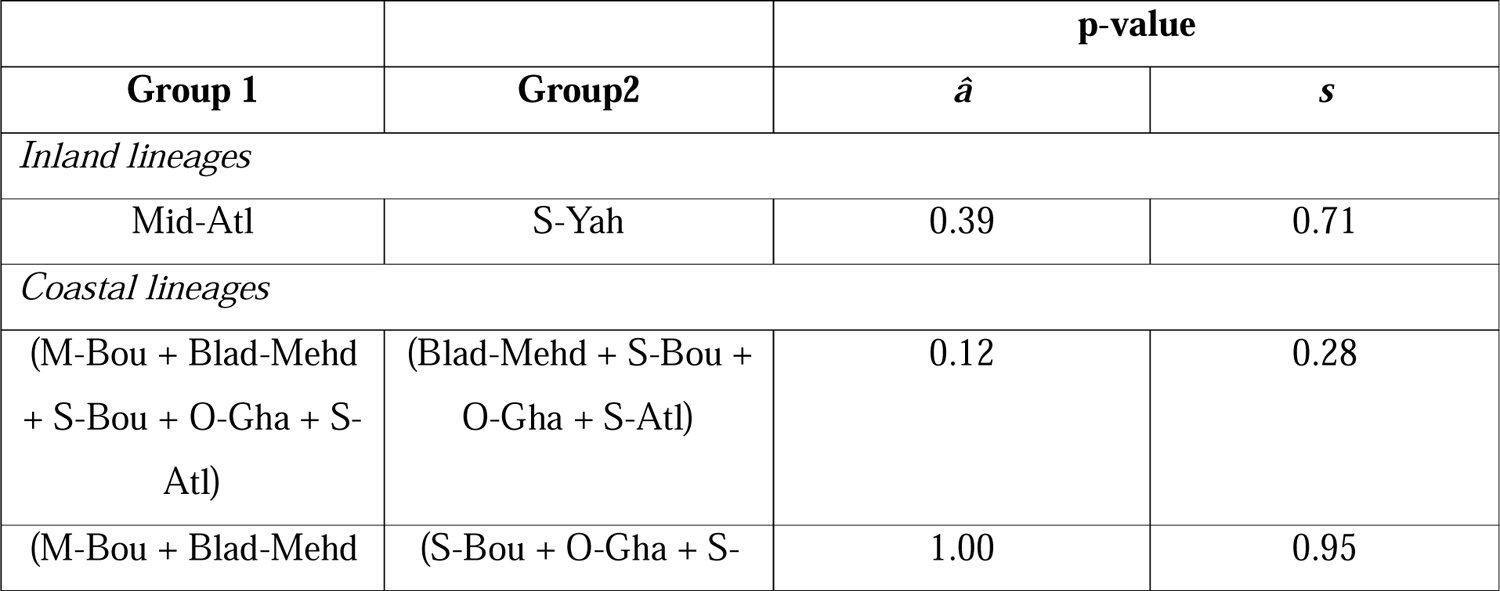
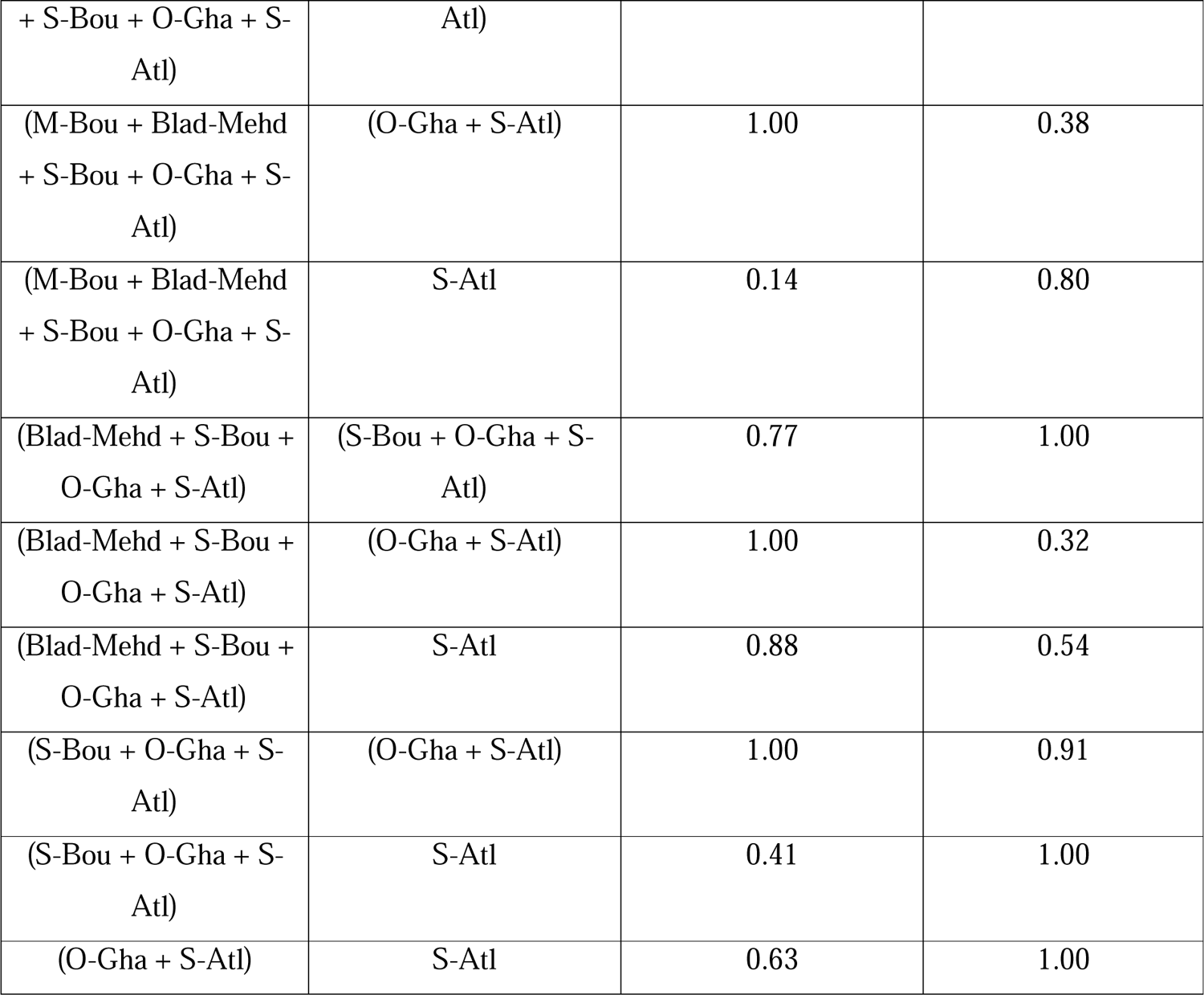
Results of simplified isolation by distance tests based on within-groups distances only, for the cases where the method of Hausdorf & Hennig (2020) could not be applied.

Starting from the north, all tests congruently supported a barrier to gene flow between N-Atl and the Rif (Fig. 3A). This was also the case when comparing Rif and Mid-Atl (Fig. 3B), to the exception of one test: based on *â* genetic distances, we couldn’t reject the null hypothesis *H032* that the between-groups regression differed from the within-group regression of Rif, although this difference was significative when considering the within-group regression of Mid-Atl (*H031*). The results for Mid-Atl and Mara (Fig. 3C) were also ambiguous: based on the *s* genetic distance, we found that the between-groups distances were different from the within-group distances of Mara, but other tests (i.e., *H02* based on *â* and *H031* based on *s*) were non-significant. On the contrary, tests between Mara and S-Atl (Fig. 3D) clearly supported a barrier to gene flow. Finally, tests based on both *â* and *s* unambiguously supported that S-Atl and Mid-Atl (Fig. 3E) differentiate along the same IBD gradient. While the differentiation between S-Yah and Mid-Atl could not be tested in the same framework because S-Yah is represented by a single locality, we were not able to distinguish regression lines calculated on the distances within Mid-Atl alone or Mid-Atl + S-Yah (Fig 3F), which we interpret as an indication that they evolve along the same IBD gradient.

On the Atlantic coast, tests based on both *â* and *s* genetic distances suggested a barrier to gene flow between N-Atl and the other coastal populations (Figure 4A, Table 2). However, these results remain inconclusive, as the regression on between-groups distances was significantly different from that on the within-group distances of N-Atl, but not the within-group distances of the rest of the coastal populations. Our tests further suggested that all coastal populations south of N-Atl differentiate along a single IBD gradient (Figure 4B).

**Figure 4.**
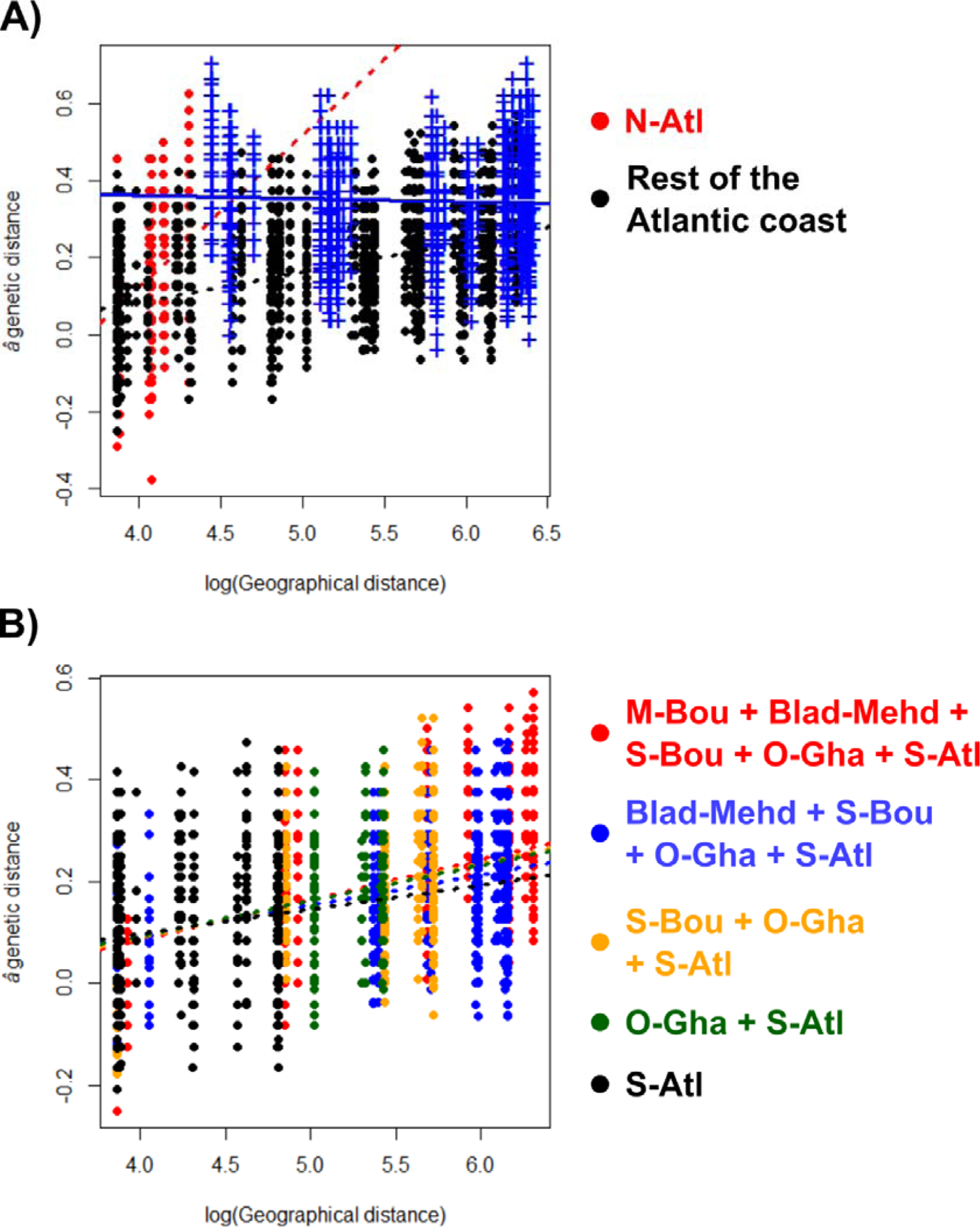
Isolation by distance tests along the Atlantic coast based on the â genetic distance. A) Comparison between N-Atl (red) and the rest of the coastal populations (black). Blue cross show between-groups distances and lines represent the respective regressions. B) Regressions on within-group distances of several combinations of coastal populations South of N-Atl.

## Discussion

### 1. Mitochondrial and nuclear markers yield contrasting phylogeographic signals in Moroccan

#### Acanthodactylus erythrurus

As already highlighted by Miralles et al. (2020), our analyses of 776 bp of the ND4 mtDNA gene supported an important genetic diversity in the Moroccan populations of *A. erythrurus*, which can be divided into several deep lineages. Our analyses recovered an origin of the Ibero-Moroccan clade (“IM” clade) around 6 Mya, a relatively old age for a species-level lineage in comparison with calibrations in *Podarcis* wall lizards (Salvi, Pinho, Mendes & Harris, 2021) and other Lacertids (Garcia-Porta et al., 2019; Joger et al., 2007; Tamar et al., 2016). Furthermore, the overall congruence between calibrations based on the end of the Messinian crisis and on fixed mutation rates seem to confirm that Iberian populations have been isolated since the re-opening of the Strait of Gibraltar, around 5.4 Mya. While the deepest nodes within the IM clade are unresolved, possibly as a result of rapid diversification, or the limited informativeness of the sequence used, we could identify 11 strongly supported (aLRT > 90) clades corresponding to geographical units (subsequently referred to as mtDNA lineages) among the Moroccan populations. While some of these mtDNA lineages include several populations distributed across relatively large areas, many well differentiated monophyletic groups corresponded to single localities, especially along the Atlantic coast. It is remarkable that in some cases geographically distant populations share closely related haplotypes (e.g., in the Mid-Atl2 and S-Atl lineages), while in other instances localities that are close to each other cluster in distinct lineages (e.g., S-Rif and Mid-Atl1 lineages meeting in Ifrane, and the localities in the Northern half of the Atlantic coast).

This pattern could suggest the existence of many phylogeographic breaks across Morocco, resulting from allopatric diversification following gene flow disruptions. Especially, in the Middle-Atlas, two highly differentiated mitochondrial lineages (S-Rif and Mid-Atl1) enter in contact with limited apparent admixture, a pattern consistent with allopatric divergence followed by secondary contact (Avise et al., 1987). Alternatively, this pattern could also result from fine-scale barriers to gene flow, including divergent ecological preferences between populations.

Contrastingly, our analyses of nine nuclear loci provided little support to the mtDNA lineages. Several “state-of-the-art” approaches, including phylogeny and STRUCTURE clustering, failed to recover consistent lineages, seemingly because of high levels of allele sharing among populations. This result was surprising, as we expected that populations evolving allopatrically for several millions of years would accumulate genome-wide mutations reflecting the mitochondrial structure. The high levels of allele sharing could thus indicate ongoing nuclear gene flow between lineages, or an important retention of ancestral polymorphisms due to large effective population sizes (Pamilo & Nei, 1988). Given the matrilineal inheritance of the mitochondria, male-biased dispersal could exacerbate differences in structure with nuclear markers (Palumbi & Baker, 1994). Finally, the poor performance of STRUCTURE to infer genetic clusters could be due to the presence of IBD, which breaks the model’s assumptions (Perez et al., 2018), resulting in over-estimating the number of genetic clusters in the data (Frantz, Cellina, Krier, Schley & Burke, 2009; Schwartz & McKelvey, 2009; Serre & Pääbo, 2004).

Contrary to other approaches based on more constrained models (i.e., phylogenetic trees, STRUCTURE), multivariate analyses of individual genotypes at the nine nuclear loci supported a phylogeographic structure somewhat consistent with the mtDNA lineages. One important difference was the position of the individuals from Sidi-Yahia (S-Yah, just south of Rabat), which harbors a very distinctive mitochondrial lineage, branching as sister to all the other populations. However, nuclear data suggest that these individuals are closely related to populations from the middle-Atlas and the Atlantic coast. Two scenarios can be considered to explain this discrepancy: firstly, the population of Sidi-Yahia could be closely related to its neighbours (i.e., Atlantic coast, Middle-Atlas), with a mitochondrial genome introgressed from a distinct genetic pool, absent from our analyses. Alternatively, Sidi-Yahia could be the remnant of an older lineage whose nuclear genome got swamped by that of invading populations from the middle-Atlas or Atlantic coast. Either way, it seems that this pattern is the legacy of an unsampled, possibly extinct *Acanthodactylus* lineage, whose mitochondrial genome persisted to that day. The existence of “ghost mitochondrial lineages” is well-documented in animals (for examples in reptiles, see Renoult, Geniez, Bacquet, Benoit & Crochet, 2009; Salvi, Pinho & Harris, 2017; Schultze et al., 2020), and their persistence can result from various mechanisms, including local adaptation (Carpio et al., 2021). Future studies of *A. erythrurus* based on genome-wide data might be useful to understand the processes underlying such discordance.

At a larger scale, in the absence of a well resolved nDNA tree, it is difficult to assess the extent of cytonuclear discordances in *A. erythrurus*. However, genetic groups identified from mtDNA and nDNA are mostly concordant, the main difference residing in the lower phylogenetic resolution yielded by nDNA markers. We hypothesize that this pattern could result from two scenarios: 1) the mitochondrial lineages reflect phylogeographic breaks, and the poor resolution of nDNA markers is the result of ancestral polymorphism retention; or 2) the mitochondrial lineages are largely connected by nuclear gene flow. We tried to untangle these processes by studying the distribution of genetic differentiation along geographical gradients.

### 2. The conundrum of separating vicariance from isolation by distance

Geographical and genetic distances were overall strongly correlated in our data, highlighting the role of IBD in generating genetic diversity in *A. erythrurus*. However, IBD alone did not explain the observed phylogeographic patterns. Using linear regressions, we could identify at least four geographic groups separated by barriers to gene flow (summarized in Figure 5, discussed in-depth in the next section), within which we could not reject the null hypothesis that populations differentiate through IBD. It is interesting to note that our results suggest that similar levels of mitochondrial divergence can arise from both IBD and allopatric divergence (and even allopatric speciation, as highlighted at the Rif/Middle-Atlas transition by Doniol-Valcroze et al., 2022). In that context, interpreting mitochondrial splits as evidence for phylogeographic breaks is potentially misleading and biased toward an over-estimation of the number of phylogeographic lineages. While contrasting mitochondrial and nuclear phylogeographic patterns can help determining the extent of mitochondrial diversity that can be attributed to vicariance, this issue remains very challenging in the absence of clear signal in nuclear markers. This problem is especially relevant as mitochondrial markers are still widely used to infer biogeographic patterns and species boundaries. In many cases, multi-locus studies employ only a single or few nuclear markers, whose signal gets eclipsed by that of the more variable mtDNA fragments. Hence, we argue that a thorough interpretation of mtDNA phylogeographic patterns should rely on a comparison with independently analyzed nuclear markers.

**Figure 5.**
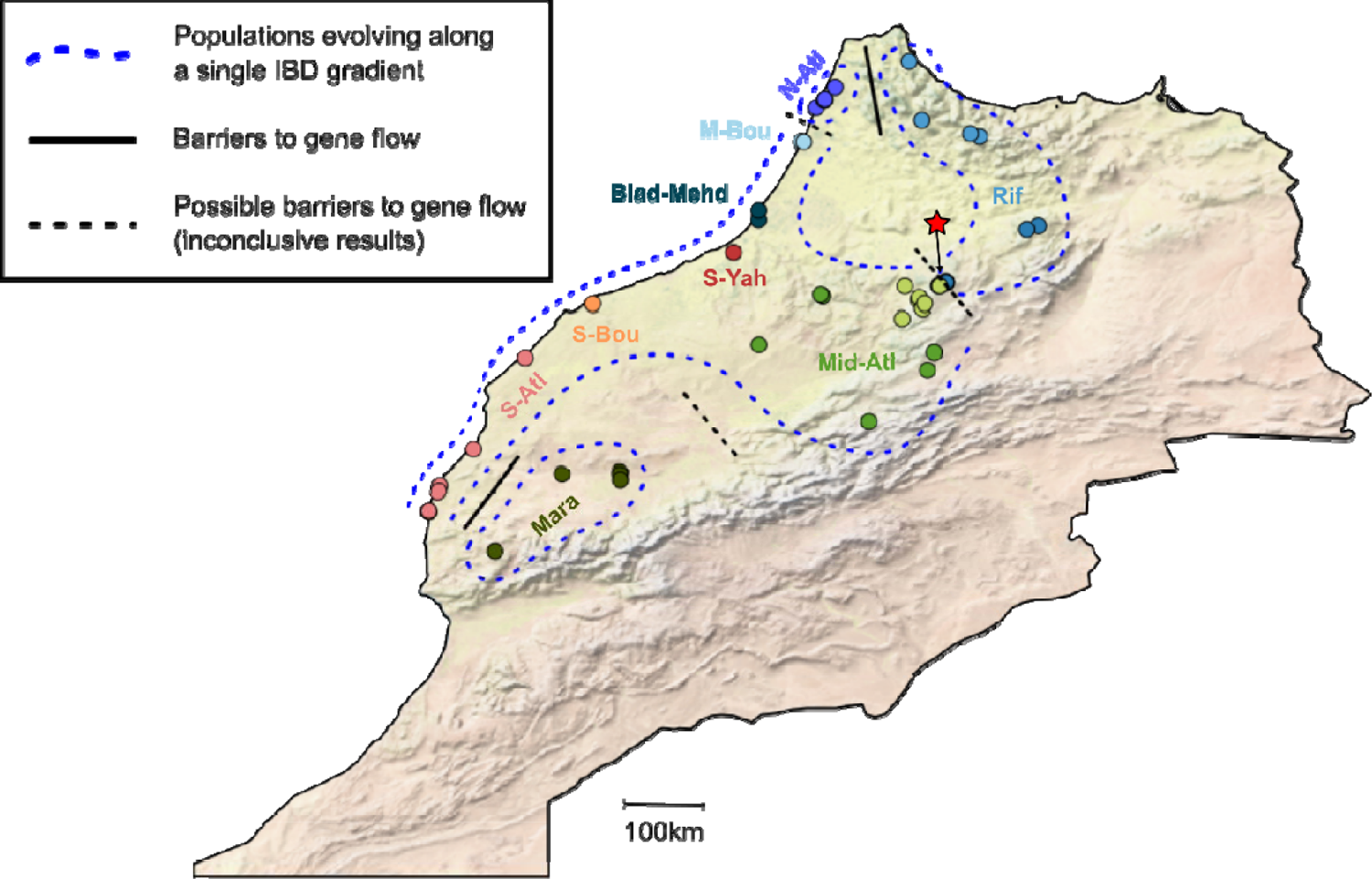
Summary of the processes underlying populations differentiation in Moroccan Acanthodactylus erythrurus. The red star highlights the contact zone between the Rif and Middle-Atlas lineages, that is inconclusive in our analysis but is identified as a species-level barrier in Doniol-Valcroze et al. (2022).

As we show here, employing statistical, spatially explicit methods to detect discontinuities in genetic diversity can yield valuable insights on the presence of phylogeographic breaks. However, our analyses often yielded inconclusive results, even in a case where we expected the signal for vicariance to be clear (i.e., the contact zone between Rif and Middle-Atlas, as inferred from the low admixture limited to a single locality, despite geographic distances being < 1km), suggesting low power of these methods even with nine independent markers. We observed a large variance in genetic distances within populations, making our tests more prone to stochastic errors. Employing more informative data (typically genome-wide markers) would likely reduce this variance, yielding more power to distinguish regression lines.

More importantly, we identify the sampling scheme as a significant factor driving the results of IBD tests. For example, it seems that the inconclusive results for the Rif/Middle-Atlas tests could be due to the asymmetric sampling around the contact zone, with only a few samples available in the former group, and a large sampling gap between the contact zone and the core area. In-depth analyses of this transition by Doniol-Valcroze et al. (2022) also identified sampling gaps as a limiting factor, even when considering genome-wide data. Likewise, our IBD tests for coastal populations were very limited because of “lineages” represented by single localities, which made thorough comparisons of within-groups and between-groups distances impossible. This issue could be mitigated by sampling many localities, especially to fill gaps between identified genetic groups.

Finally, the IBD model used assumes that genetic and geographic distances follow a linear relationship (Hausdorf & Hennig, 2020). Because most genetic distance metrics are comprised between 0 and 1, this assumption is likely to be broken when genetic distances are high. Especially, intra- and between-groups regressions can be difficult to distinguish when intra-group distances are close to 1 (Hausdorf & Hennig, 2020). However, the linear relationship is expected at least in one case, when considering log-transformed geographic distances and *Fst*/(1-*Fst*) as an estimate of genetic distance (and, by extension, the *â* genetic distance for individuals; Rousset, 1997). In our case, intra-group distances were very variable and sometimes very large, especially in S-Atl and Mid-Atl, which might result in false negatives or inconclusive results. Considering more markers (e.g., genome-wide SNPs) might improve this issue by reducing the variability across individuals and avoiding very high distances. Furthermore, violations of the linearity assumption might explain the differences in results between the *â* and *s* genetic distances.

To conclude, our results demonstrate that identifying phylogeographic breaks is far from a trivial issue and should not rely on mitochondrial markers alone. It would be interesting to further study the factors underlying the concordance of mitochondrial breaks with vicariance events, especially the effects of dispersal abilities and population sizes.

### 3. Mechanisms of diversification in the Acanthodactylus erythrurus group

Small terrestrial vertebrates, including lizards, often have low dispersal abilities, which sometimes result in strong phylogeographic patterns (e.g., Alfonso Silva et al., 2017; Potter et al., 2019). Hence, it is not surprising to find a high genetic diversity associated with strong IBD gradients in *A. erythrurus*. However, low dispersal alone could not explain the phylogeographic pattern observed in *A. erythrurus* across Morocco, and we found at least four groups separated by barriers to gene flow (Figure 5): the northern Atlantic Coast, the Rif, the surroundings of Marrakech, and a large group including the remaining the Atlantic coast, the Middle-Atlas and Sidi-Yahia. However, several of the inferred barriers to gene flow were not unambiguously supported, highlighting a need for further investigations. Surprisingly, this was the case for Rif and Middle-Atlas, two groups that show very little to no traces of admixture at their contact zone, near Ifrane, where individuals belonging to either group occur 1 km apart. An in-depth analysis of this contact zone based on genome-wide data by Doniol-Valcroze et al. (2022) showed a very steep transition, demonstrating that the Middle-Atlas and Rif are strongly differentiated groups with restricted admixture, consistent with the existence of intrinsic barriers to gene flow (Dufresnes, Pribille, et al., 2020; see Doniol-Valcroze et al. (2022) for a more in-depth discussion of the Rif/Middle-Atlas transition). This result supports the hypothesis that ancestral polymorphism retention in our nine loci dataset, rather than widespread gene flow, blurred the phylogeographic signal, to the point that two lineages at an advanced stage of speciation were sometimes difficult to separate. Both the northern Atlantic and Marrakech plains populations showed patterns of divergence akin that of the Rif (i.e., monophyly in the nDNA tree, clear divergence in the MCAs), and might be candidates for deep intra-specific - or even species-level - lineages. Future studies employing a larger number of molecular markers will hopefully help clarifying how these groups are related, and whether they are linked by extensive gene flow or not.

Although the application of our approach was limited by lineages represented by single localities, our results support that populations from the Middle-Atlas, Atlantic coast to the exclusion of the three northernmost localities, and Sidi-Yahia are all part of the same IBD gradient, without barriers to gene flow detected. Alternatively, it is also possible that the Middle-Atlas and Atlantic coast represent two formerly allopatric lineages that resumed high levels of gene flow upon secondary contact, as supported by the intermediate position of some coastal populations (especially Blad-Dendoun and Mehdia) in both MCAs and population trees. In the absence of explicit measures of admixture, such scenario would be difficult to distinguish from strict IBD. Interestingly, these populations are separated into two morphologically well differentiated subspecies, *A. e. atlanticus* (Middle-Atlas) and *A. e. lineomaculatus* (Atlantic coast), the latter having even be proposed as a distinct species (Bons & Geniez, 1995). Our results suggest that this morphological variation could have arisen, or been maintained, in the face of gene flow, maybe as a result of local adaptation.

Contrary to what we observed in *A. erythrurus*, phylogeographic studies of reptile species widespread in Western and Northern Morocco mostly report low levels of genetic diversity within Morocco (e.g., El-Farhati et al., 2021; Faria & Harris, 2020; Nicolas et al., 2018; Rosado, Rato, Salvi & Harris, 2017; but see Klesser et al., 2021 and Veríssimo et al., 2016 for more complex patterns). Some of the phylogeographic breaks we identified here correspond with large rivers (e.g., Oued Loukos between N-Atl and the other costal Atlantic populations), others with ecological transitions (soft coastal grounds versus hard inland substrates between S-Atl and Mara) but most do not align with obvious landscape or ecological breaks. It is possible that a substantial fraction of the lineages we identified correspond to different species (as for the Rif and Middle-Atlas lineages; Doniol-Valcroze et al., 2022), and that their divergences are maintained by intrinsic barriers to gene flow, similarly to the various species of the wall lizards (*Podarcis*) that currently inhabit the Iberian Peninsula (Caeiro-Dias et al., 2018, Caeiro-Dias, Rocha et al., 2021; Caeiro-Dias, Brelsford et al., 2021). Further sampling around contact zones and analyses of gene flow across these contact zones with genomic markers will be needed to partition the effects of intrinsic (e.g., mate recognition, selection against hybrids) and extrinsic (geography and ecology) barriers to gene flow in *A. erythrurus* in North Africa.

Whatever the current mechanisms underlying phylogeographic breaks in the species in Morocco, it is hard to pinpoint precisely which processes generated such a diversity over a relatively small spatial scale. It is possible that this pattern is the legacy of populations fragmentation caused by climatic variations during the Pleistocene, when periods of high aridity occurred (Le Houérou, 1997). *Acanthodactylus erythrurus* populations, being associated with Mediterranean and mesic habitats (Fonseca et al., 2009), might have repeatedly retracted to cooler and more humid places during arid periods, such as mountain ranges or coasts (similar to “refugia-within-refugia”, Gómez & Lunt, 2007). Our results highlight that the recent biogeographic history of taxa within Morocco, and the Maghreb in general, is still poorly understood, and call for more comparative phylogeography studies in this area, especially relying on multi-locus data sets.

## Acknowledgements

The authors are grateful to Raouaa Fathalla and Patricia Sourrouille for their help with laboratory work, and to Christian Hennig for his helpful input on linear models. Part of the analyses were carried out by L.R. using resources in project 2022/22-653 provided by the Swedish National Infrastructure for Computing (SNIC) at UPPMAX, partially funded by the Swedish Research Council through grant agreement no. 2018-05973. Part of this work was carried out by using the resources of the national INRAe MIGALE (Migale bioinformatics Facility, https://doi.org/10.15454/1.5572390655343293E12) and GENOTOUL (Bioinfo Genotoul, https://doi.org/10.15454/1.5572369328961167E12) bioinformatics HPC platforms, as well as the local Montpellier Bioinformatics Biodiversity (MBB, supported by the LabEx CeMEB ANR-10-LABX-04-01) and CBGP HPC platform services. This research was conducted in the scope of the LIA “Biodiversity and Evolution”.

## Biosketch

**Loïs Rancilhac** is an early-career evolutionary biologist broadly interested in the genetics and genomics of speciation. This work started as his master thesis and was further developed as a secondary project during his PhD. It is part of a broader project focused on the consequences of low dispersal and speciation on the genetic structure of *Acanthodactylys erythrurus*, led by Pierre-André Crochet and Raphaël Leblois in Montpellier, France.

## Authors contributions

A.M., J.C.B., M.B., P.A.C., P.G. and R.L. collected samples. P.A.C. and P.G. managed the samples. A.M., M.B. and D.M.A. generated the genetic data. A.M. performed preliminary explorations of the dataset. P.A.C, L.R. and R.L. conceived the research idea. L.R. performed the analyses presented in the manuscript with contributions from A.M., P.A.C and R.L. L.R. wrote the manuscript with contributions from P.A.C. All authors revised the manuscript.

