## Supplementary materials for "Phylogeographic breaks and how to find them: An empirical attempt at separating vicariance from isolation by distance in a lizard with restricted dispersal"

| **Number** | **Name** | **Longitude** | **Latitude** | **Sample size** |
| --- | --- | --- | --- | --- |
| 1 | Souk Tnine (SKTN) | 35.3971 | -5.9661 | 2 |
| 2 | Souk Khemis_1 (SKEM) | 35.2794 | -6.0664 | 12 |
|  | Souk Khemis_2 (SKEM) | 35.2848 | -6.0621 | 13 |
| 3 | Larache (LARA) | 35.2086 | -6.1418 | 21 |
| 4 | Martil (MART) | 35.6379 | -5.2776 | 6 |
| 5 | Bab Taza (TAZA) | 35.0899 | -5.1595 | 9 |
| 6 | Ketama_1 (KETA) | 34.9373 | -4.6158 | 10 |
|  | Ketama_2 (KETA) | 34.9647 | -4.70559 | 5 |
| 7 | Tazzeka_1 (TZKA) | 34.1042 | -4.0724 | 1 |
|  | Tazzeka_2 (TZKA) | 34.0708 | -4.1806 | 1 |
| 8 | Ifrane-North (IFRA-N) | 33.5828 | -4.9352 | 3 |
| 9 | Moulay Bousselham_1 (MBOU) | 34.896 | -6.2877 | 14 |
|  | Moulay Bousselham_2 (MBOU) | 34.8878 | -6.2583 | 1 |
| 10 | Mehdia (MEHD) | 34.2485 | -6.6798 | 10 |
| 11 | Blad Dendoun (BLAD) | 34.15902 | -6.68007 | 7 |
| 12 | Ifrane-South_1 (IFRA-S) | 33.5732 | -4.9315 | 11 |
|  | Ifrane-South_2 (IFRA-S) | 33.577 | -4.9349 | 2 |
|  | Ifrane-South_3 (IFRA-S) | 33,5449 | -5,0004 | 1 |
|  | Ifrane-South_4 (IFRA-S) | 33.5433 | -4.9927 | 1 |
| 13 | Azrou_1 (AZRU) | 33.4145 | -5.1919 | 1 |
|  | Azrou_2 (AZRU) | 33.43516 | -5.18188 | 6 |
|  | Azrou_3 (AZRU) | 33.5427 | -5.3169 | 13 |
|  | Azrou_4 (AZRU) | 33.33586 | -5.15701 | 3 |
|  | Azrou_5 (AZRU) | 33.3818 | -5.13239 | 1 |
|  | Azrou_6 (AZRU) | 33.24586 | -5.34916 | 4 |
| 14 | Col Zad (CZAD) | 32.9297 | -5.0465 | 1 |
| 15 | Boumia (BOUM) | 32.7655 | -5.1081 | 22 |
| 16 | Ouaourioud (OUAO) | 32.2939 | -5.6593 | 1 |
| 17 | Oulmès_1 (OULM) | 33.4497 | -6.0812 | 12 |
|  | Oulmès_2 (OULM) | 33.4507 | -6.0835 | 5 |
|  | Oulmès_3 (OULM) | 33.463 | -6.102 | 2 |
| 18 | Oued Zem (OZEM) | 32.9995 | -6.6789 | 1 |
| 19 | Sidi Yahia (SYAH) | 33.8553 | -6.9129 | 24 |
| 20 | Sidi Boumaine (SBOU) | 33.3798 | -8.2261 | 10 |
| 21 | Marrakech_1 (MARA) | 31.8176 | -7.9784 | 10 |
|  | Marrakech_2 (MARA) | 31.7921 | -7.9812 | 2 |
|  | Marrakech_3 (MARA) | 31.7389 | -7.9783 | 1 |
| 22 | Sidi Chiker (SDCH) | 31.7965 | -8.5215 | 9 |
| 23 | Timlite (TIMT) | 31.0883 | -9.1391 | 3 |
| 24 | Ouled Ghanem (OGHA) | 32.8792 | -8.859 | 7 |
| 25 | Souira (SOUI) | 32.032 | -9.3406 | 22 |
| 26 | Essaouira_1 (ESSA) | 31.6869 | -9.6632 | 2 |
|  | Essaouira_2 (ESSA) | 31.636 | -9.6738 | 3 |
|  | Essaouira_3 (ESSA) | 31.4582 | -9.7595 | 6 |
|  | Essaouira_4 (ESSA) | 31.4545 | -9.756 | 18 |
|  | Essaouira_5 (ESSA) | 31.4636 | -9.7563 | 6 |

**Table S1.** Details of the sampled localities. The first column refers to the numbers used on Figure 1 and Table 1. On several occasions, multiple nearby sampling localities were lumped under the same name, but coordinates and sample sizes are given for each individually.

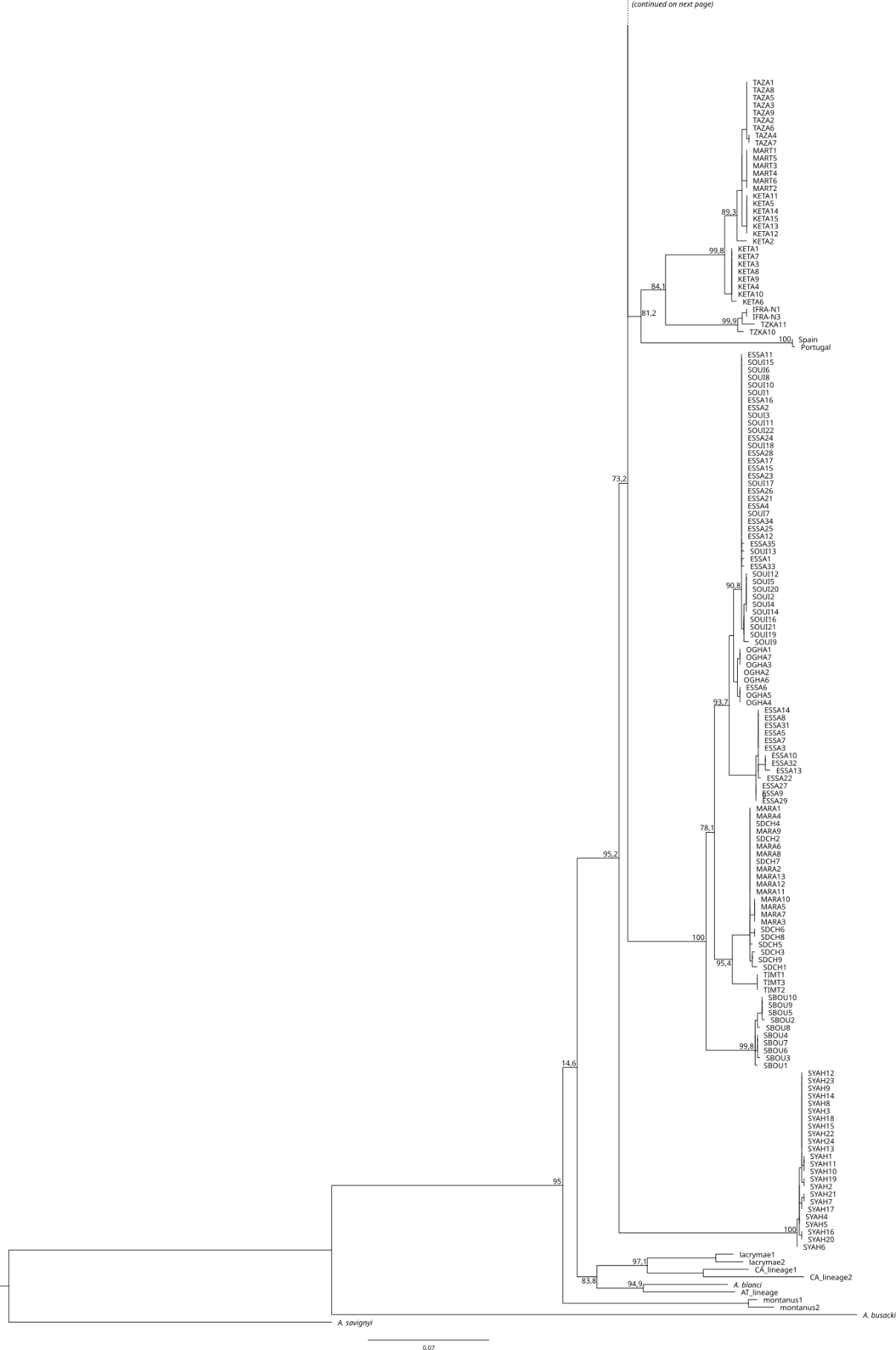

**Figure S1.** Complete mitochondrial tree inferred from 776 bp of the ND4 gene. Samples names refer to the localities listed in Table S1. Approximate-Likelihood ratio test support of the nodes is given (not represented for the shallowest nodes for improved clarity). (Continued on next page)

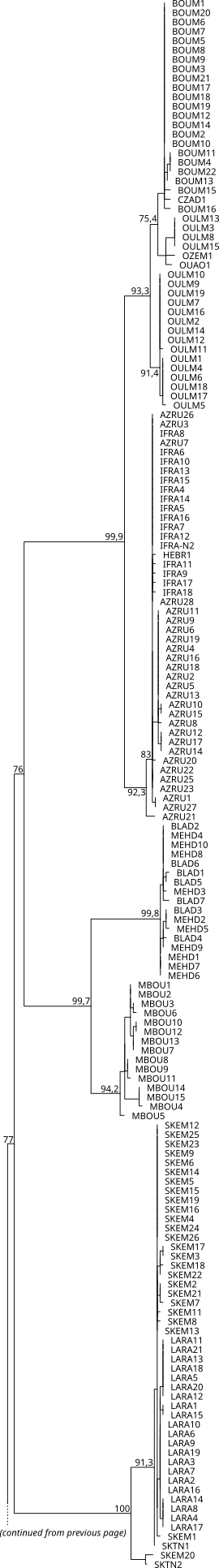

**Supplementary S1: Calibration of the mitochondrial phylogeny**

We used 776 bp of the ND4 mitochondrial gene to estimate divergence times between lineages of *A. erythrurus* under two approaches. First, we constrained the split between the Iberian populations and their closest Moroccan relatives at 5.4 Mya [ 5.3, 5.5 ], corresponding to the end of the Messinian salinity crisis, during which land bridged the South of the Iberian peninsula and North Africa. Secondly, to avoid possible bias from post-Messinian oversea dispersal, we also calibrated the tree using a fixed mutation rate. Since no direct estimation of the mutation rate (μ) of the ND4 mitochondrial gene exist for Lacertidae, we extrapolated it from published data on the Cytochrome b (CytB) gene. To do so, we first downloaded sequences of CytB and ND4 genes from Genbank for 21 Lacertidae species (Table S2). We then plotted pairwise p-distances on ND4 as a function of pairwise p-distances on CytB for all species pairs. We found that both distances followed a significant positive correlation ( p < 0.01) with a coefficient of 0.93. Hence, we considered that $\mu\left( ND4 \right)=0.93\times\mu(CytB)$. No direct estimation of µ(CytB) exist for *Acanthodactylus* species, but it has been estimated in *Podarcis* (µ=0.013 substitutions/site/My, Brown et al. 2008) and *Gallotia* (µ=0.015 substitutions/site/My, Cox et al. 2010). Hence, we calibrated the tree twice, with both mutation rates.

The analyses were conducted in BEAST v. 2.6.7 (Bouckaert et al. 2019) under a strict molecular clock and a GTR+Γ4 substitution model (corresponding to the closest model available to the best fitting model found by ModelFinder). Preliminary runs showed that these analyses were computationally intensive and failed to reach convergence even with very long chains of 500,000,000 steps. Hence, we subsampled our dataset to 18 sequences (one per mitochondrial lineage), and for each analysis we performed three independent runs (manually setting different seeds for each) of 10,000,000 steps, sampling every 5,000. The results of the three runs were combined in LogCombiner v. 2.6.3 (Bouckaert et al. 2019) with the first 30% steps discarded as burnin. A Maximum Clade Credibility tree was built from the posterior distribution in TreeAnnotator v. 2.6.3 (Bouckaert et al. 2019). Even with this approach, the analysis did not reach proper convergence. However, since we do not use the inferred dates for downstream inferences but just to get an idea of the timescale of divergences, we argue that this approach should be sufficient.

**Cited literature:**

Bouckaert, R., Vaughan, T. G., Barido-Sottani, J., Duchêne, S., Fourment, M., Gavryushkina, A., Heled, J., Jones, G., Kühnert, D., De Maio, N., Matschiner, M., Mendes, F. K., Müller, N. F., Ogilvie, H. A., du Plessis, L., Popinga, A., Rambaut, A., Rasmussen, D., Siveroni, I., Suchard, M. A., Wu, C.-H., Xie, D., Zhang, C., Stadler, T., & Drummond, A. J. (2019). BEAST 2.5: An advanced software platform for Bayesian evolutionary analysis. *PLoS computational biology*, *15*(4), e1006650.

Brown, R. P., Terrasa, B., Pérez-Mellado, V., Castro, J. A., Hoskisson, P. A., Picornell, A., & Ramon, M. M. (2008). Bayesian estimation of post-Messinian divergence times in Balearic Island lizards. *Molecular Phylogenetics and Evolution*, *48*(1), 350-358.

Cox, S. C., Carranza, S., & Brown, R. P. (2010). Divergence times and colonization of the Canary Islands by *Gallotia* lizards. *Molecular Phylogenetics and Evolution*, *56*(2), 747-757.

**Table S2.** Accession numbers of the CytB and ND4 sequences of 21 Lacertidae species used to calculate the relationship between the mutation rates of the two genes.

| **Species** | **Accession number CytB** | **Accession number ND4** |
| --- | --- | --- |
| *Algyroides fitzingeri* | GQ142134.1 | KX081004.1 |
| *Algyroides marchi* | GQ142133.1 | KX081005.1 |
| *Algyroides moreoticus* | GQ142131.1 | KX081001.1 |
| *Algyroides nigropunctatus* | GQ142132.1 | KX080999.1 |
| *Dalmatolacerta oxycephala* | GQ142129.1 | KX081049.1 |
| *Dinarolacerta montenegrina* | GQ142141.1 | KX081010.1 |
| *Dinarolacerta mosorensis* | GQ142130.1 | KX081015.1 |
| *Hellenolacerta graeca* | GQ142128.1 | KX081050.1 |
| *Iberolacerta monticola* | GQ142124.1 | KX081023.1 |
| *Iranolacerta brandtii* | GQ142140.1 | KX081061.1 |
| *Lacerta agilis* | GQ142118.1 | KX081040.1 |
| *Lacerta bilineata* | AY714981.1 | KX081043.1 |
| *Lacerta dugesii* | GQ142121.1 | KX081036.1 |
| *Phoenicolacerta kulzeri* | GQ142139.1 | KX081059.1 |
| *Podarcis muralis* | JF422107.1 | KX081018.1 |
| *Podarcis siculus* | AY185095.1 | KX081016.1 |
| *Psammodromus algirus* | DQ150367.1 | KX081069.1 |
| *Scelarcis perspicillata* | GQ142122.1 | KX081031.1 |
| *Takydromus sexlineatus* | GQ142142.1 | KX081052.1 |
| *Timon lepidus* | GQ142119.1 | KX081030.1 |
| *Zootoca vivipara* | GQ142120.1 | KX081033.1 |

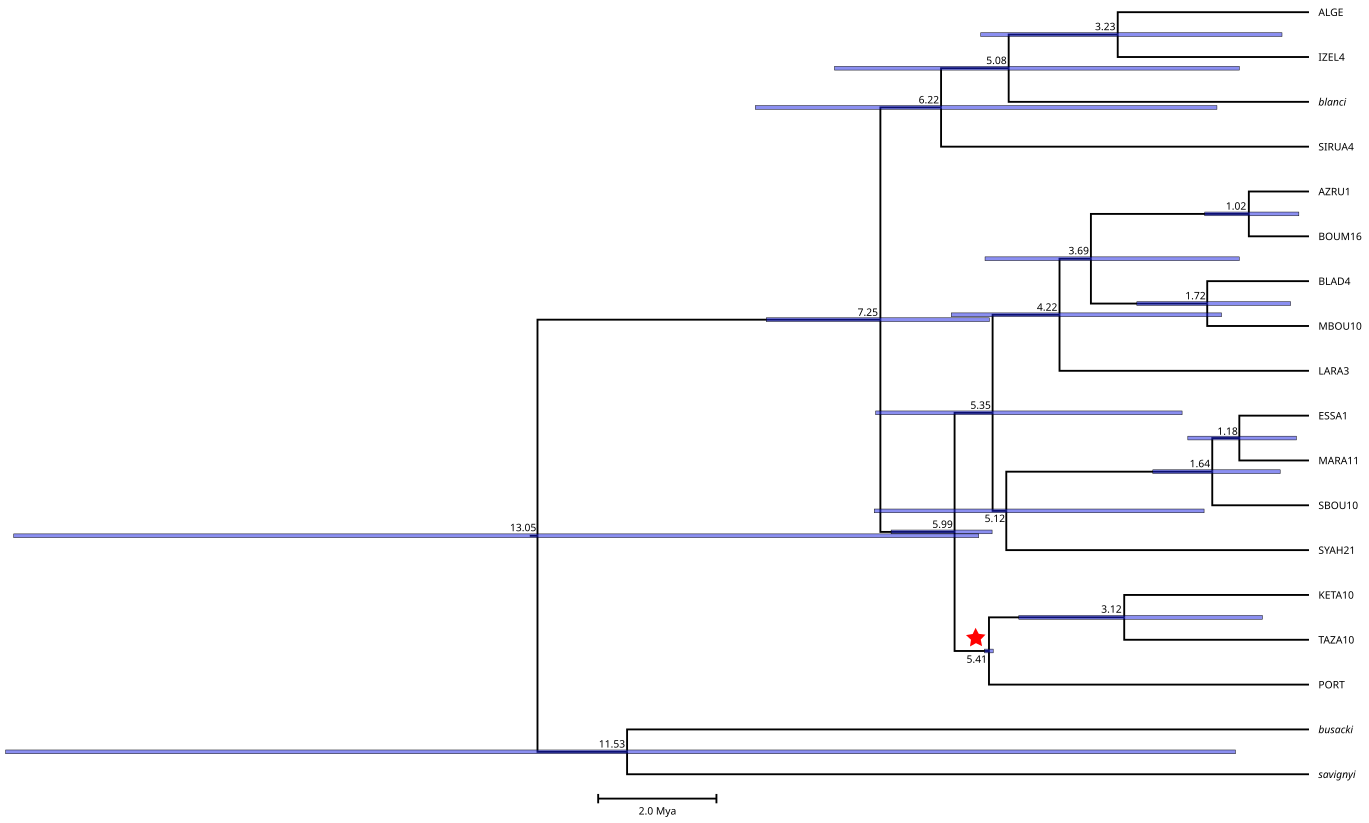

**Figure S2.** Time-tree estimated from 18 sequences of the ND4 mitochondrial gene under a strict molecular clock, using the end of the Messinian crisis as a calibration for the divergence between the Iberian populations and their closest Moroccan relatives (node marked with a red star). The numbers at the nodes indicate their average age, and the blue bars show 95% confidence intervals.

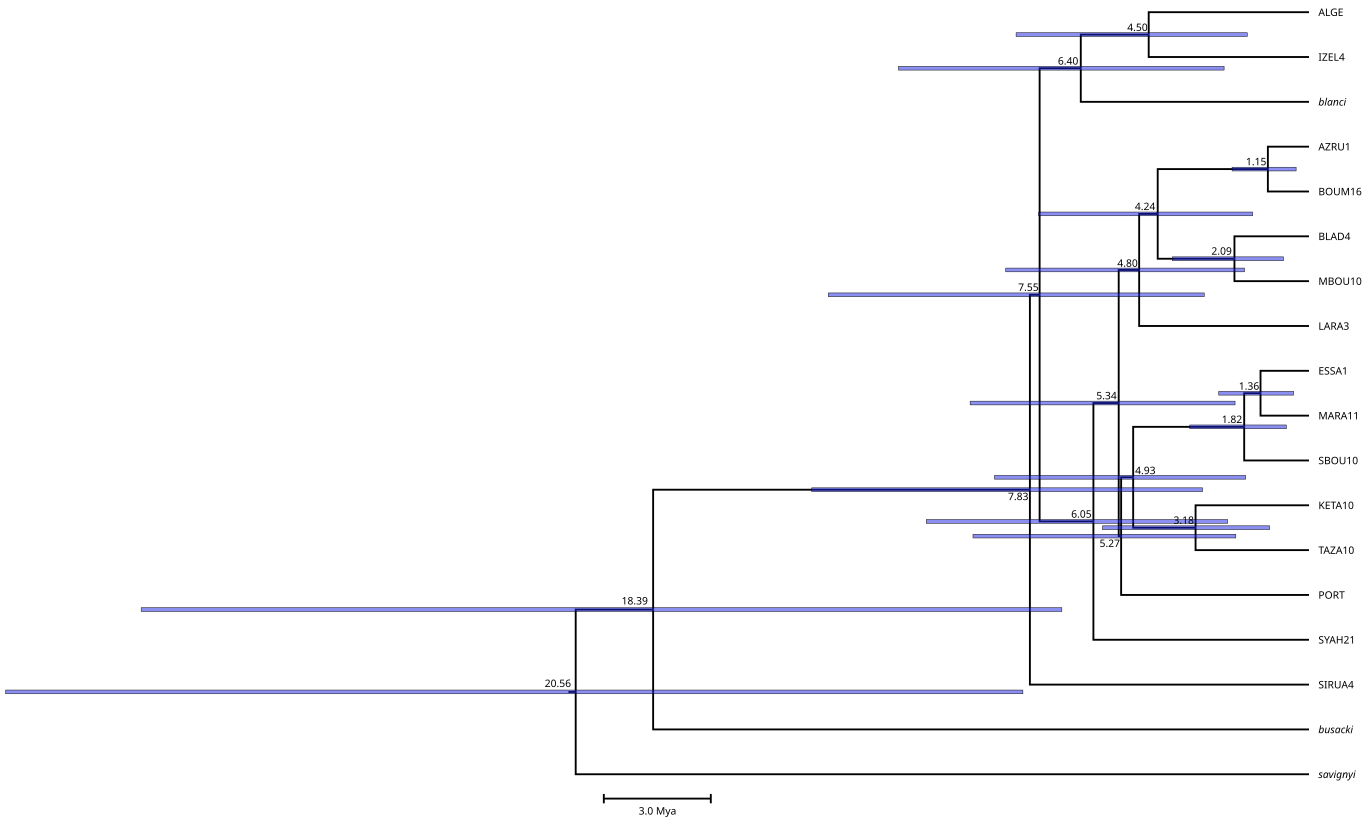

**Figure S3.** Time-tree estimated from 18 sequences of the ND4 mitochondrial gene under a strict molecular clock, using a fixed mutation rate estimated from the Cytochrome B mutation rate of *Podarcis*. The numbers at the nodes indicate their average age, and the blue bars show 95% confidence intervals.

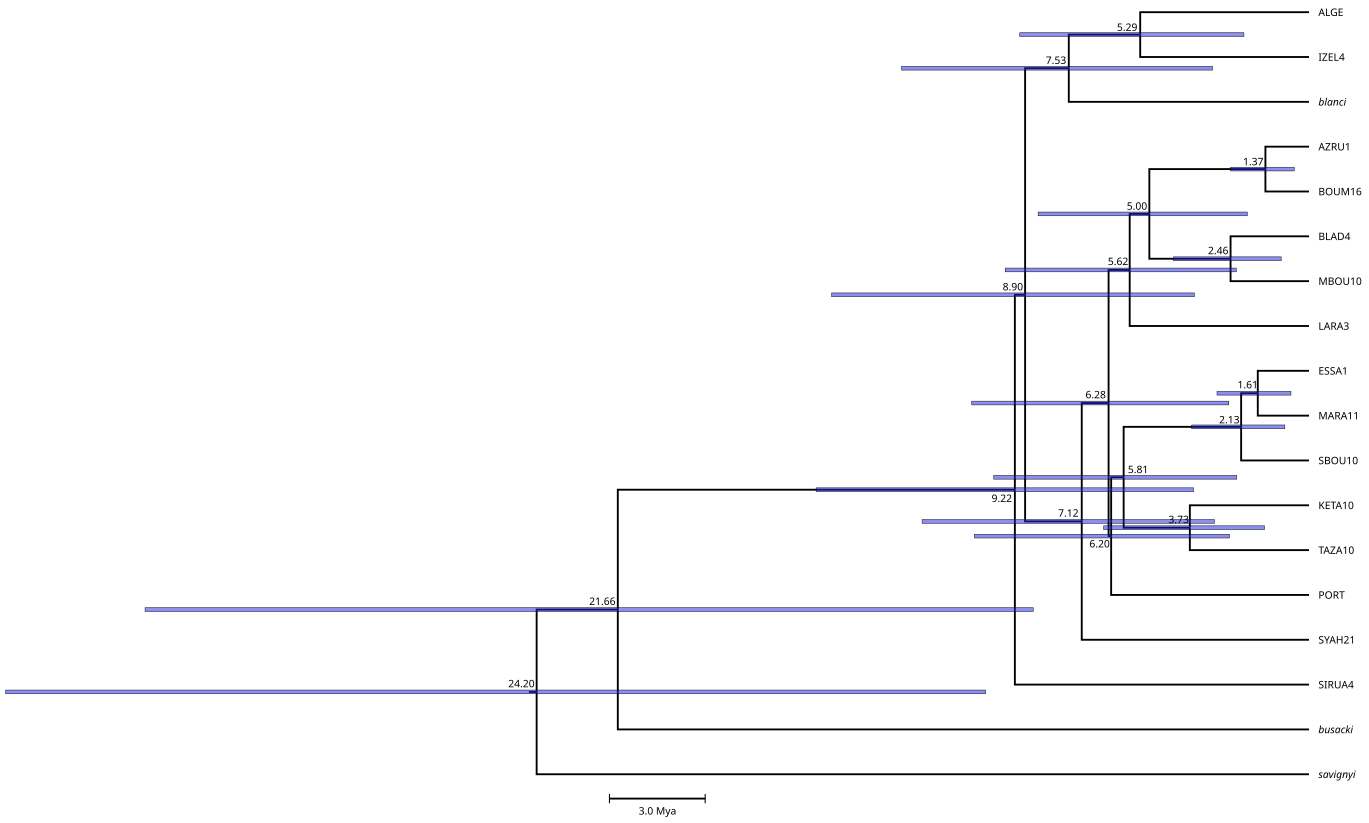

**Figure S4.** Time-tree estimated from 18 sequences of the ND4 mitochondrial gene under a strict molecular clock, using a fixed mutation rate estimated from the Cytochrome B mutation rate of *Gallotia*. The numbers at the nodes indicate their average age, and the blue bars show 95% confidence intervals.

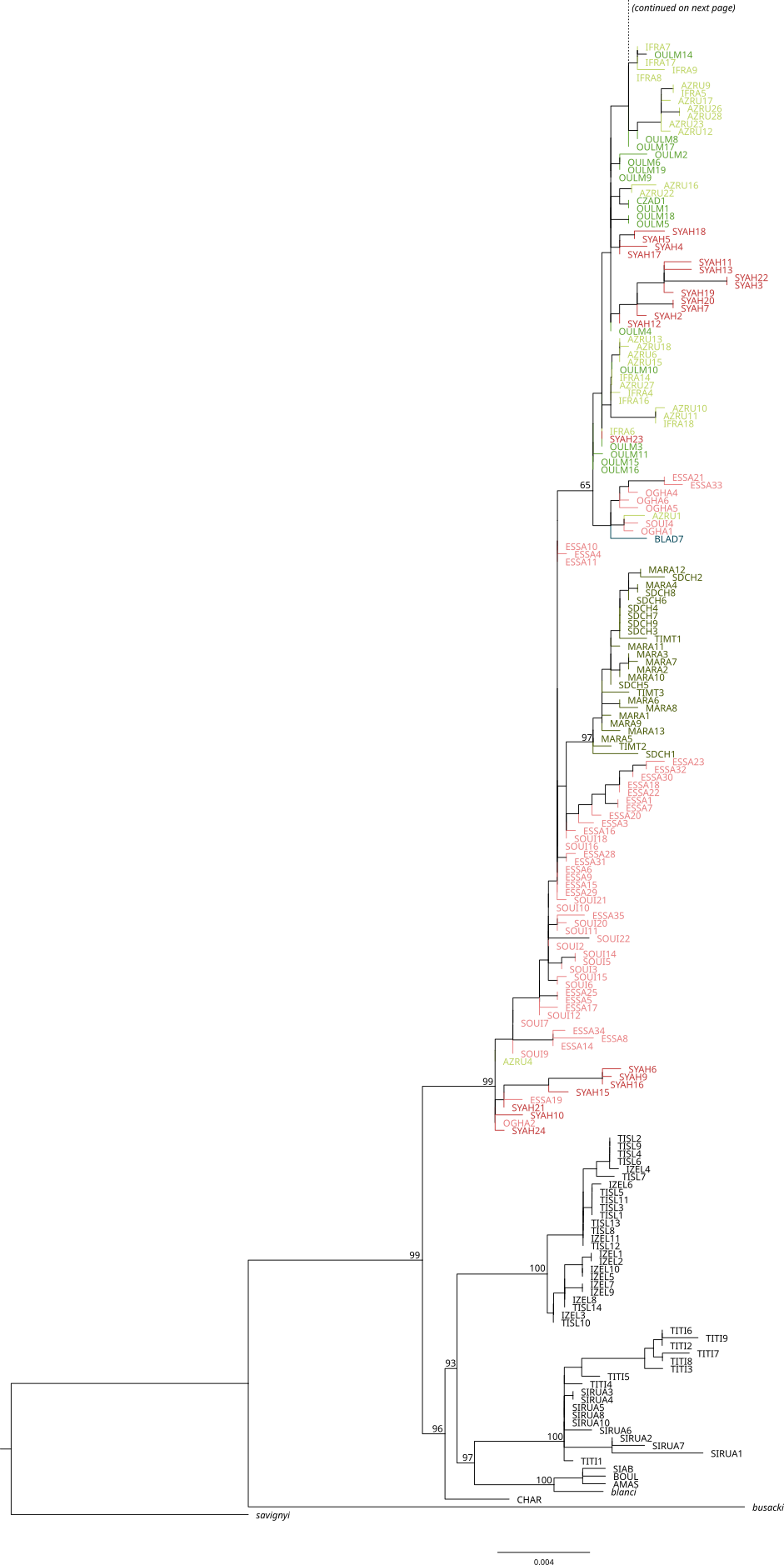

**Figure S5.** Maximum-likelihood tree inferred from the concatenation of the nine nuclear introns. Approximate Likelihood Ration Test supports are given for the internal nodes (not represented at shallow nodes for improved clarity). Terminal branches and samples names are colored based on the mitochondrial lineage they belong to (black branches are outgroups), following the color code of Figure 1. (Continued on next page)

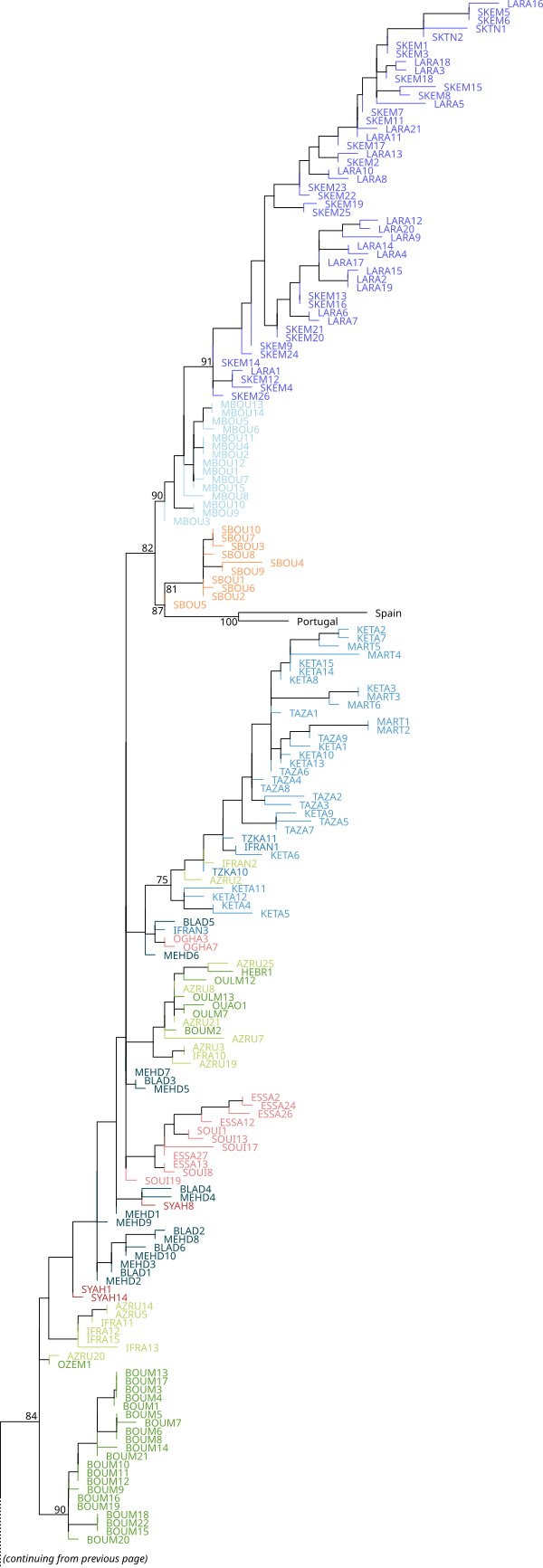

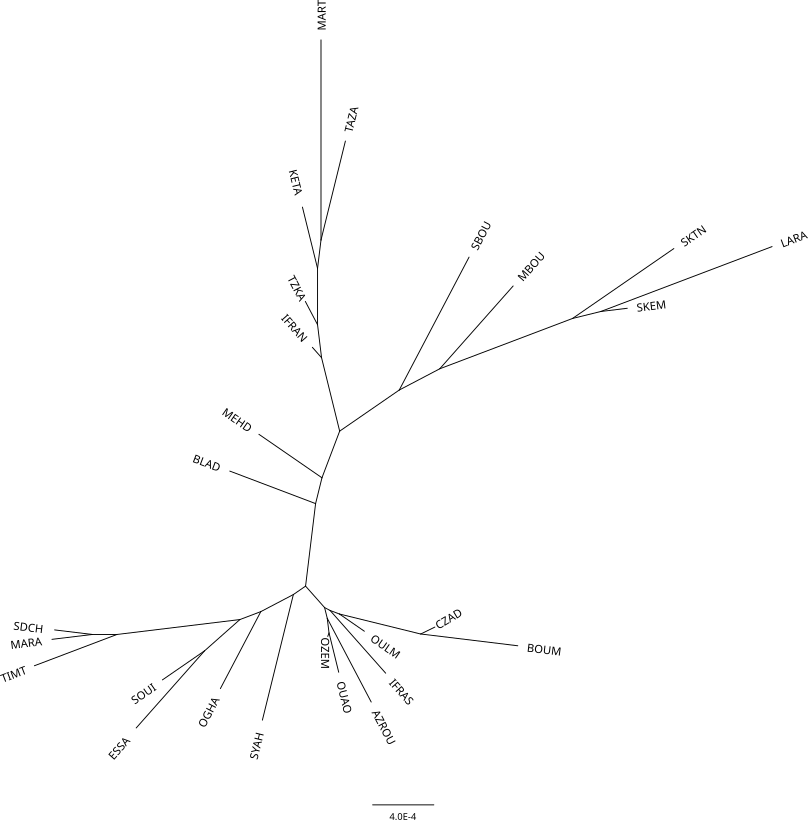
**Figure S4.** Neighbor-Joining tree calculated from the average p-distances between populations.

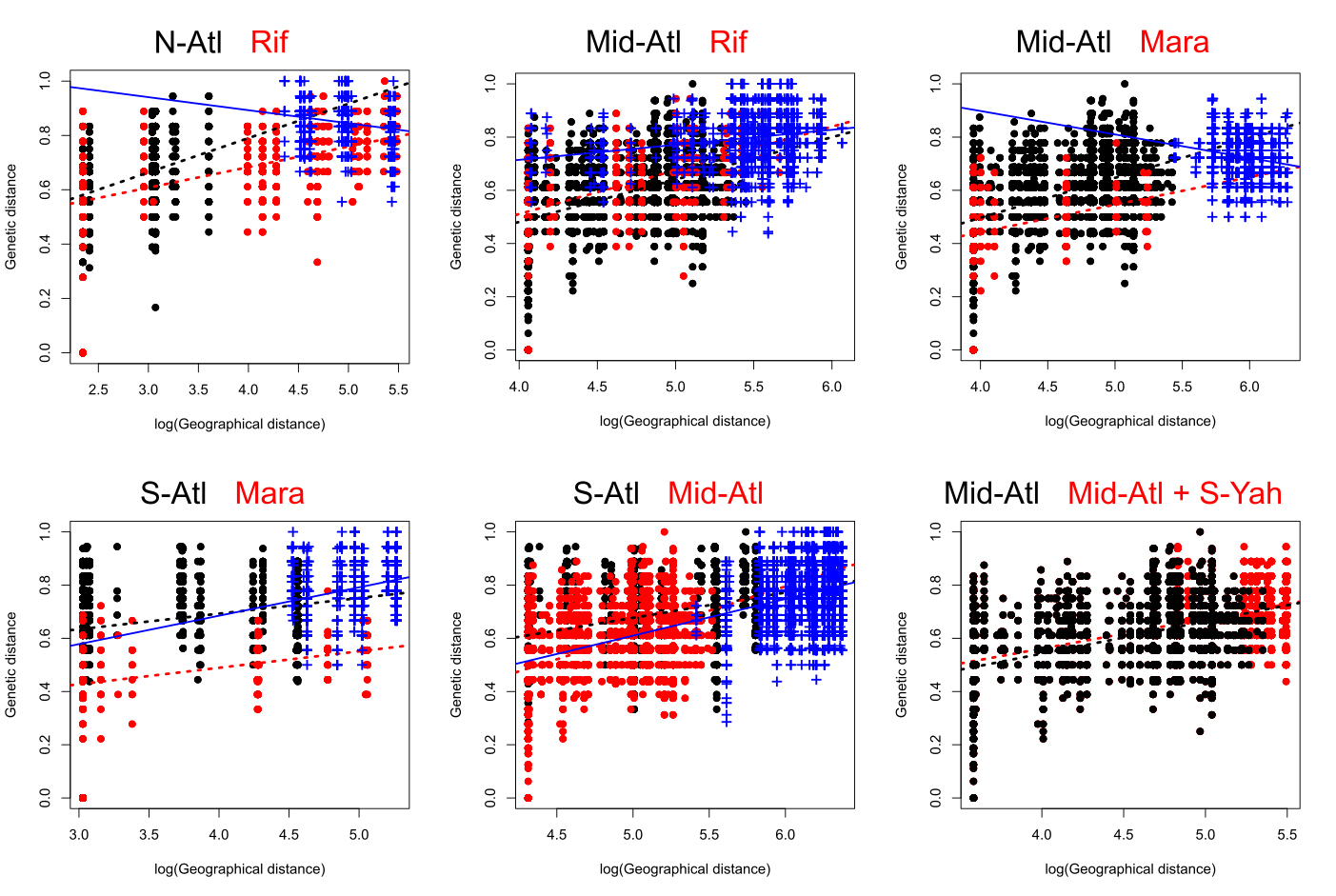

**Figure S7.** Linear regressions between genetic (s) and geographical distances in pairs of geographically adjacent inland lineages. Black and red dots show within-group distances for both lineages, and dashed lines are linear regression fitted to them. Blue crosses and solid lines show between-group distances. Between-group distances are not represented on the last graph as the test was adapted to account for zero-only geographical distances in S-Yah.

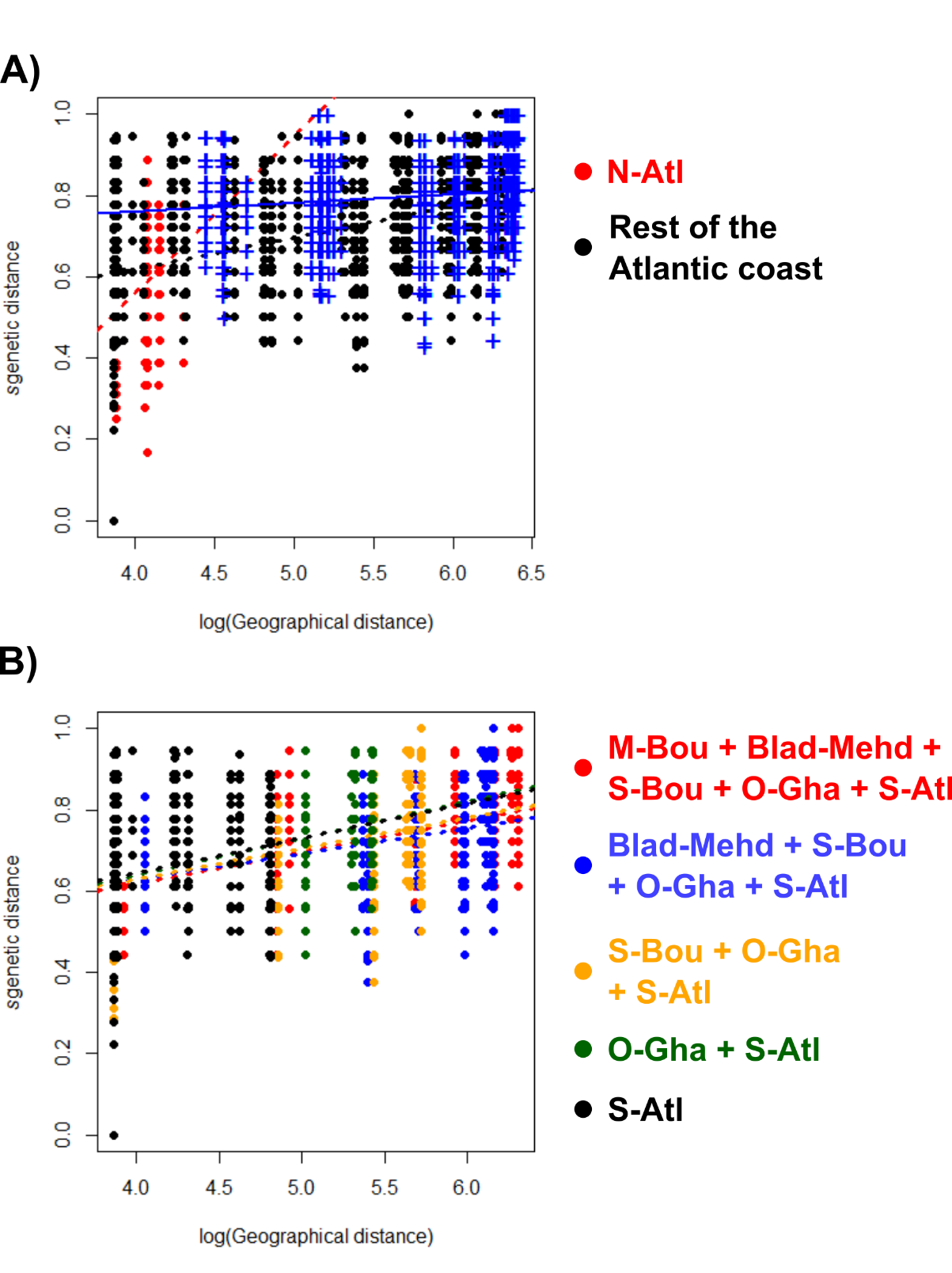

**Figure S8.** IBD tests along the Atlantic coast based on the shared alleles genetic distance. A) Comparison between N-Atl (red) and the rest of the coastal populations (black). Blue cross show between-groups distances and lines represent the respective regressions. B) Regressions on within-group distances of several combinations of coastal populations South of N-Atl.
